# The epigenetic landscape of oligodendrocyte progenitors changes with time

**DOI:** 10.1101/2024.02.06.579145

**Authors:** David K. Dansu, Sami Sauma, Dennis Huang, Meng Li, Sarah Moyon, Patrizia Casaccia

## Abstract

**SUMMARY:** Dansu et al. identify distinct histone H4 modifications as potential mechanism underlying the functional differences between adult and neonatal progenitors. While H4K8ac favors the expression of differentiation genes, their expression is halted by H4K20me3.

Adult oligodendrocyte progenitors (aOPCs) generate myelinating oligodendrocytes, like neonatal progenitors (nOPCs), but they also display unique functional features. Here, using RNA-sequencing, unbiased histone proteomics analysis and ChIP-sequencing, we define the transcripts and histone marks underlying the unique properties of aOPCs. We describe the lower proliferative capacity and higher levels of expression of oligodendrocyte specific genes in aOPCs compared to nOPCs, as well as the greater levels of H4 histone marks. We also report increased occupancy of the H4K8ac mark at chromatin locations corresponding to oligodendrocyte-specific transcription factors and lipid metabolism genes. Pharmacological inhibition of H4K8ac deposition reduces the levels of these transcripts in aOPCs, rendering their transcriptome more similar to nOPCs. The repressive H4K20me3 mark is also higher in aOPCs compared to nOPCs and pharmacological inhibition of its deposition results in increased levels of genes related to the mature oligodendrocyte state. Overall, this study identifies two histone marks which are important for the unique transcriptional and functional identity of aOPCs.

## INTRODUCTION

Adult oligodendrocyte progenitors (aOPCs) are slowly proliferative cells, distributed throughout the gray and white matter and representing 5-8% of the total adult brain cells (Dawson et al., 2003). These cells possess electrical signaling properties and have been shown to receive neuronal synapses (Bergles et al., 2000; Gallo et al., 2008; S. C. Lin & Bergles, 2004) and retain the ability to form new myelin upon neuronal stimulation (Gibson et al., 2014; Ortiz et al., 2019). For instance, aOPCs differentiate in response to motor learning and other cognitive tasks (Bacmeister et al., 2020; McKenzie et al., 2014; Pan et al., 2020) and generate new myelinating oligodendrocytes following demyelination (Shen et al., 2008; Sim et al., 2002). Adult OPCs share with neonatal oligodendrocyte progenitors (nOPCs) the immunoreactivity for progenitor markers, although they differ in terms of proliferation and migration (Wolswijk et al., 1990; Wolswijk & Noble, 1989) and response to external signal (Lin et al., 2009). It is thought that aOPCs derive from nOPCs which escaped the differentiation program characteristic of developmental myelination. We and others previously highlighted the importance of transcription factors (Emery & Lu, 2015; Hernandez & Casaccia, 2015; Pruvost & Moyon, 2021) and histone and DNA modifications responsible for the differentiation of nOPCs into oligodendrocytes during developmental myelination (Dansu et al., 2020; Selcen et al., 2023). For instance, it was shown that proliferating nOPCs are characterized by euchromatic nuclei, enriched with acetylated lysine residues in the tail of histone H3 and that the transition from growth arrest to differentiation is initiated by repressive deacetylation, mediated by HDACs (Marin-Husstege et al., 2002; Shen et al., 2005) and responsible for the downregulation of transcriptional inhibitors of differentiation (Magri et al., 2014; Swiss et al., 2011; Wu et al., 2012). The deposition of methyl groups on specific lysine residues of Histone 3, mediated by SUV39H1/2 and EZH2 was also shown to be critical for the repression and downregulation of genes related to alternative lineages, cell cycle and electrical properties (Boshans et al., 2019; Liu et al., 2015). Additional epigenetic mechanisms of gene repression such as DNMT1-mediated DNA methylation (Moyon et al., 2016) and recruitment of repressive heterochromatin to the nuclear lamina (Pruvost et al., 2023) were shown to be important for the acquisition and maintenance of the differentiated oligodendrocyte phenotype, which also required DNA hydroxymethylation (Moyon et al., 2021) and ATP-dependent SWI/SNF chromatin-remodeling complexes (Yu et al., 2013). Overall, this suggests that specific epigenetic marks define the path leading nOPCs towards mature oligodendrocytes during developmental myelination.

In this study, we hypothesized that by escaping differentiation during developmental myelination, aOPCs may adopt histone marks that are distinct from those used by nOPCs and that this underlying epigenetic signature may account for the functional differences between the two cell states. Using RNA-seq, unbiased histone proteomics and ChIP-sequencing, we identify and characterize histone marks and target genes which contribute to defining the transcriptional and functional identity of aOPCs.

## RESULTS

### The transcriptome of nOPCs is distinct from that of aOPCs

To begin characterizing the intrinsic differences between nOPCs and aOPCs, we reanalyzed our RNA-sequencing data of cells sorted from *Pdgfra-H2BEGFP* reporter mice (Hamilton et al., 2003) at postnatal day 2 (P2) and 60 (P60) respectively **(Fig. 1A).** Principal component analysis (PCA) plots showed a clear separation between the nOPC and aOPC samples **(Supplemental Fig. S1A)** and differential gene expression analysis revealed 3,406 genes significantly upregulated and 5,082 genes significantly downregulated in aOPCs, using a log_2_FoldChange cut off of >0.75 and FDR cut-off of <0.05 **(Fig. 1A-B).** Among the down-regulated genes we noted several cell cycle genes, including cyclins regulating the G1/S transition, such as cyclin D1 and D2, encoded by *Ccnd1* and *Ccnd2* **(Fig. 1B),** while among the upregulated genes we detected several transcripts characteristic of the differentiated state, including transcription factors (e.g. *Nkx6.2*), myelin proteins (e.g. *Cnp, Mog*) and enzymes characteristic of mature oligodendrocytes (e.g. *Aspa, Ptdgs*). Indeed, heat map representations of transcripts upregulated in aOPCs identified several genes related to lipid metabolism (**Fig. 1C**), while those with decreased expression were mostly related to cell cycle and proliferation (**Fig. 1D**). Consistently, DAVID gene ontology analysis of the upregulated genes in aOPCs identified biological processes related to lipid metabolic process, regulation of transcription from RNA pol II (**Supplemental Fig. S1B),** as well as autophagy and ubiquitination **(Supplemental Fig. S1C),** consistent with recently published proteomic data in aOPCs (de la Fuente et al., 2020). In addition, biological processes such as cell adhesion, cell cycle and cell migration were identified among the signature of genes downregulated in aOPCs **(Supplemental Fig. S1D),** and were better visualized using heat maps for specific transcripts corresponding to cell adhesion and extracellular matrix **(Supplemental Fig. S1D),** consistent with previously published transcriptomic datasets (Neumann et al., 2019; Spitzer et al., 2019). Together, these results further highlight functional divergence reflected by the transcriptomic differences between nOPCs and aOPCs.

**Figure 1.**
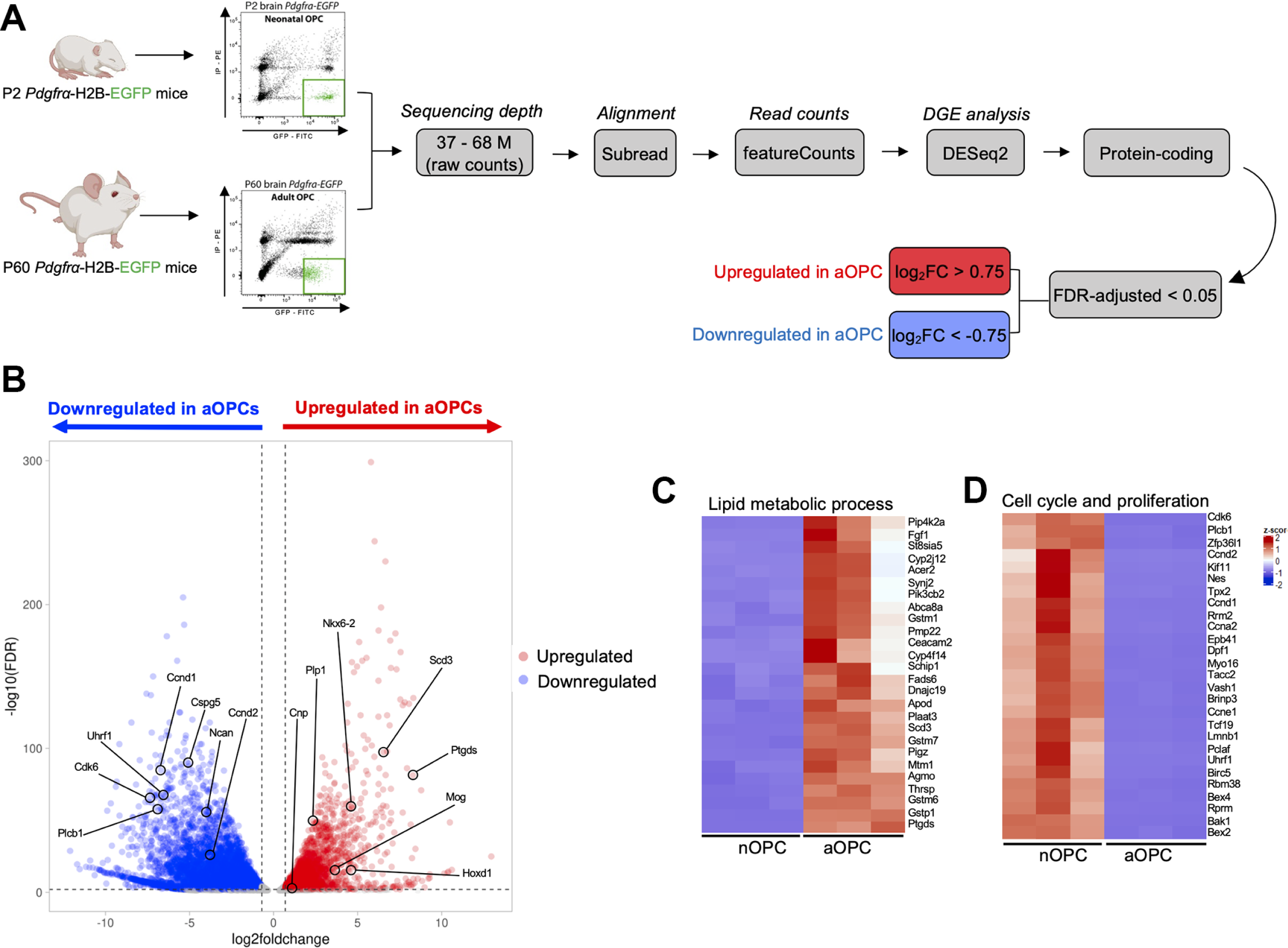
The transcriptome of adult oligodendrocyte progenitors is distinct from that of the neonatal progenitors. **(A).** Flowchart of the experimental sequence, from cell sorting of nOPCs and aOPCs from Pdgfra-H2BEGFP reporter mice to RNA-sequencing analysis. **(B).** Volcano plot showing the differentially expressed genes in aOPCs relative to nOPCs. Differentially downregulated transcripts in aOPCs are shown in blue and differentially upregulated genes are shown in red. **(C-D).** Gene expression heatmaps of selected biological processes including upregulated transcripts in aOPC related to lipid metabolic process **(C)** and down-regulated transcripts in aOPC, related to cell cycle and proliferation **(D).**

The RNA-sequencing analysis revealed higher levels of oligodendrocyte specific transcripts in aOPCs compared to nOPCs, which were validated by RT-qPCR. They included *Cnp* - encoding for the myelin protein 2’,3’-Cyclic nucleotide 3’ phosphodiesterase, *Mog*-encoding for myelin oligodendrocyte glycoprotein (**Fig. 2A)** and *Nkx6-2* (Cai et al., 2010; Southwood et al., 2004), encoding for the homeobox transcription factor NKX6.2 (**Fig. 2B)** and *Ptgds (*Pan et al., 2023), encoding for the Prostaglandin D2 Synthase enzyme **(Fig. 2C)**. In order to rule out that these findings could be due to the potential contamination of the aOPC samples with mature oligodendrocytes, we used RNAScope fluorescent *in situ* hybridization and a probe specific for *Pdgfra* in order to unequivocally identify progenitors in the adult or neonatal brain. The detection of this marker and of myelin transcripts such as *Cnp* **(Fig. 2D)** or the enzyme *Ptgds* **(Fig. 2E)** only in sections from the adult (P60) and not neonatal P5 brain, further supported the expression of these oligodendrocyte specific markers in aOPCs.

**Figure 2.**
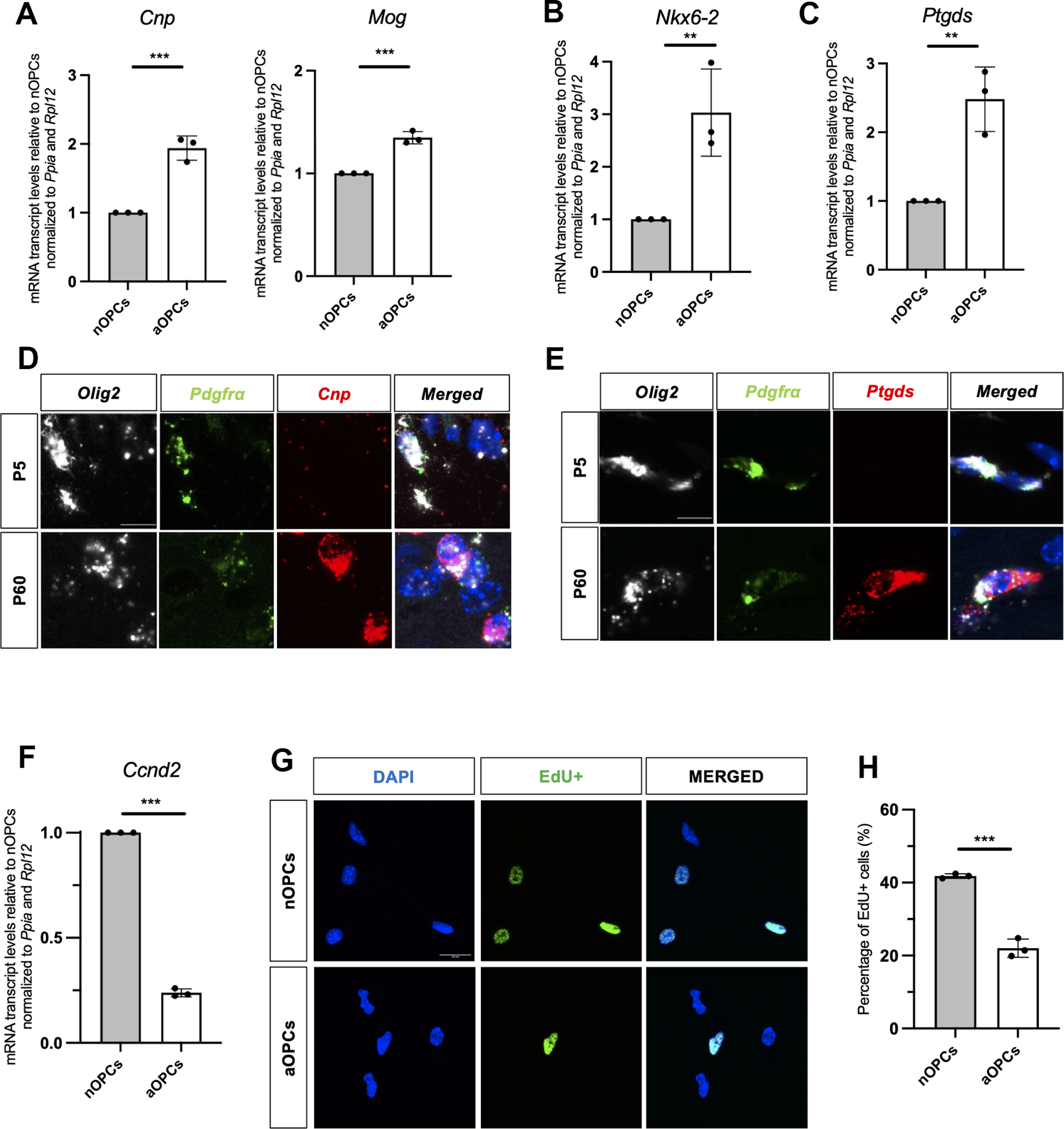
Higher transcript levels of oligodendrocyte specific genes and lower transcript levels of genes related to proliferation in adult compared to neonatal OPCs. **(A-C)**. Quantitative real-time PCR validation of selected transcripts (Cnp, Mog, Nkx6-2 and Ptgds) which were upregulated in aOPCs (white bars) compared to nOPCs (gray bars). Data represent average values relative to nOPC samples ± SD for n = 3 (**p <0.01, ***p <0.001, Student’s t test). **(D-E).** In situ RNA scope hybridization results showing the transcript levels of Cnp (D) and Ptgds (E) in *Olig2*^+^ (white) and *Pdgfra*^+^ (green) cell in P5 and P60 coronal mice brain sections. Cnp and Ptgds transcripts are shown in red. DAPI is shown in blue. Scale bar = 10 μm. **(F).** Quantitative real-time PCR validation of the levels of Ccnd2 in aOPCs (white bars) compared to nOPCs (gray bars). Data represent average values normalized to nOPC samples ± SD for n = 3 (***p <0.001, Student’s t test). **(G).** Representative confocal image of nOPCs and aOPCs after 5 h of EdU incorporation for assessment of cell proliferation. DAPI^+^ and EdU^+^ cells shown in blue and green respectively. Scale bar = 50 μm. **(H).** Quantification of percentage of EdU+ cells in nOPCs (gray bars) and aOPCs (white bars). Data represent average values normalized to nOPC samples ± SD for n = 3 (***p<0.001, Student’s t test).

Besides expressing higher levels of differentiation genes, aOPCs were also characterized by lower levels of cell cycle related transcript, Cyclin D2 *(Ccnd2)*, as measured by RT-qPCR and compared to nOPCs **(Fig. 2E).** Incorporation of 5-ethynyl-2’-deoxyuridine (EdU) over five hours was then used to assess the proliferative rate of cultured aOPCs and nOPCs kept in the same experimental conditions. Consistent with previous reports (Wolswijk et al., 1990; Wolswijk & Noble, 1989), we detected a smaller percentage of EdU incorporating aOPCs (x= 22.0% ± 2.5%, n = 274 cells counted from 3 biological experiments) compared to nOPCs (x =41.8% ± 0.6% n = 204 cells counted from 3 biological experiments) **(Fig. 2G, H)**. Together, these data validated the existence of transcriptional and functional differences between aOPCs and nOPCs and led us to investigate the epigenetic landscape of OPCs with age.

### Identification of histone post-translational modifications distinguishing aOPCs and nOPCs

To begin addressing the question of age-induced epigenetic differences occurring in OPCs, we performed an unbiased proteomic analysis of the post-translational modifications of nucleosomal histones extracted from the nuclei of nOPCs and aOPCs, sorted from the brain of *Pdgfra-H2B-EGFP* reporter mice **(Fig. 3A)**. The analysis identified several histone modifications of lysine residues in the tails of histones H3 and H4 **(Supplemental Fig. S2A-B).** While several modifications of histone H3 (e.g. H3K27ac, H3K27me3) were not consistently detected and validated as different between the two cell populations (**Figure 3B-D and Supplemental Table 1**), the levels of several post-translational modifications of residues in histone H4 were consistently found to be differentially abundant in aOPCs compared to nOPCs **(Fig. 3B, and Supplemental Table 1)** and were further validated, using western blot analysis. Among those, we identified higher levels of the activating mark H4K8 acetylation (H4K8ac) **(Fig. 3F)** and of the repressive trimethylation of lysine 20 (H4K20me3) **(Fig. 3G)**. Importantly, not all residues in histone H4 were characterized by higher levels of PTMs in aOPCs compared to nOPCs, as, for instance H4K16ac levels did not differ between the two populations **(Fig. 3F).** Also, not all the lysine residues in histone H4 showed higher levels of acetylation, as we detected deacetylation of lysine 5 (H4K5ac), previously related to differentiation of nOPCs into oligodendrocytes (Scaglione et al., 2018) **(Fig. 3G).** The differential levels of H4K5ac and H4K8ac detected in sorted nuclei were further validated by immunohistochemistry (IHC) of brain sections from neonatal and adult *Pdgfra-H2BEGFP* reporter mice, using antibodies specific to H4K5ac and H4K8ac. Consistent with the histone proteomic and western blot analyses, immunohistochemistry of the corpus callosum **(Supplemental Fig. S2C-F)** and of the cortex **(Fig. 4)**, revealed lower intensity for H4K5ac **(Fig. 4A-B)** and higher intensity for H4K8ac in a OPCs compared to nOPCs **(Fig. 4C-D)**. Taken together, these results identify specific post-translational modifications of lysine residues in histone H4 as more prevalent in aOPCs than in nOPCs and suggest that they may contribute to the transcriptional and functional differences between the two cell types.

**Figure 3.**
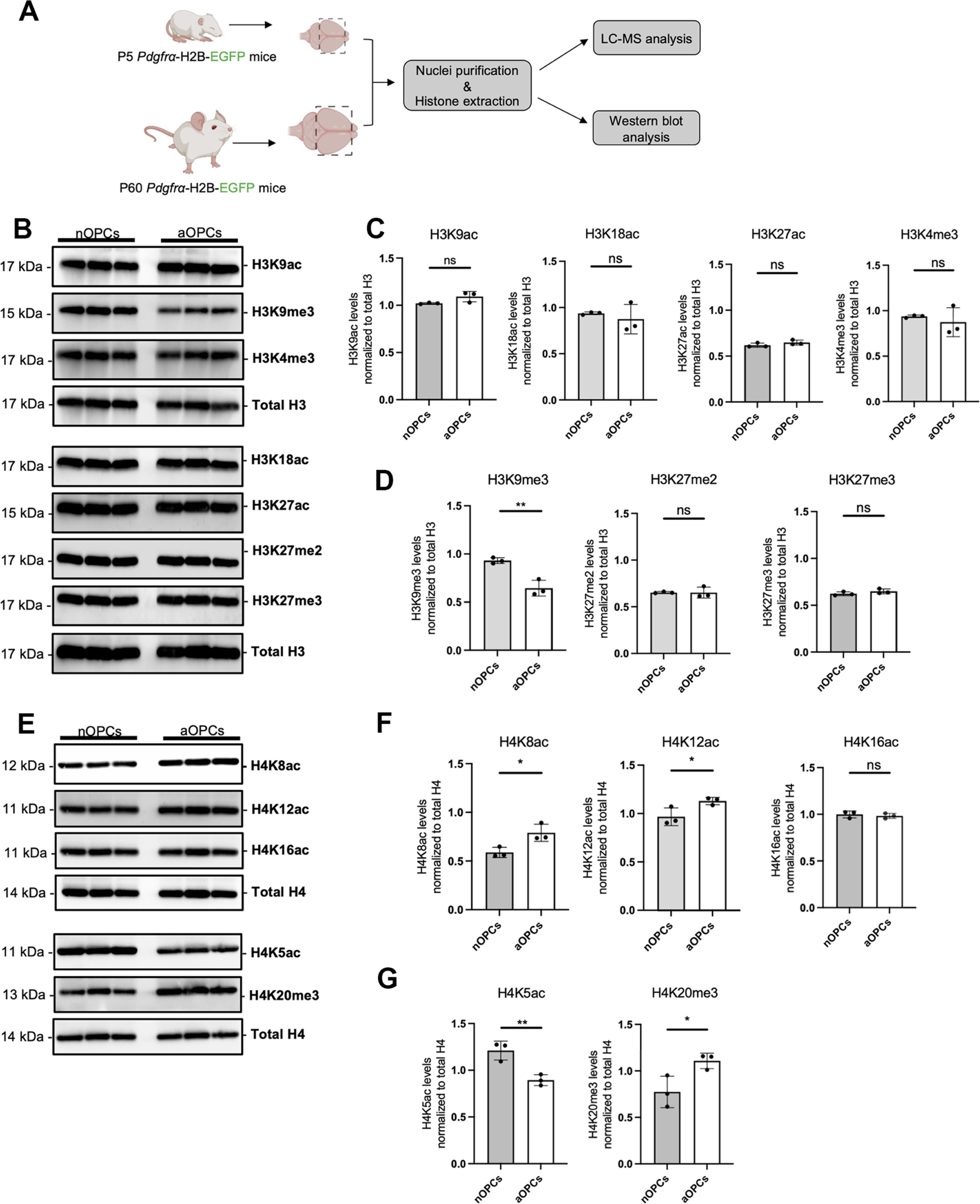
Adult oligodendrocyte progenitors are characterized by distinct histone marks compared to neonatal progenitors. **(A).** Flowchart of the unbiased quantitative histone proteomic analysis performed on extracted histones from nOPCs and aOPCs isolated from Pdgfra-H2BEGFP reporter mice. **(B).** Western blots of histone extracts from nOPCs and aOPCs, documenting the levels of the indicated post-translational modification of lysine residues in histone H3, using specific antibodies. **(C-D)**. Quantification of each of the activating H3 marks (H3K9ac, H3K18ac, H3K27ac, H3K4me3) in panel **(C)** and of the repressive marks (H3K9me3, H3K27me2 and H3K27me3) levels in panel **(D).** Data represent average values normalized to total histone H3 ± SD for n = 3 (**p < 0.01 Student’s t test). **(E)**. Western blots of histone extracts from nOPCs and aOPCs, documenting the levels of the indicated post-translational modification of lysine residues in histone H4. **(F-G)**. Quantification of activating marks (H4K8ac, H4K12ac, H4K16ac) in panel **F** and of the repressive marks (low H4K5ac and H4K20me3) in panel **G**. Data represent average values normalized to total histone H4 ± SD for n = 3 (*p < 0.05, **p < 0.01 Student’s t test).

**Figure 4.**
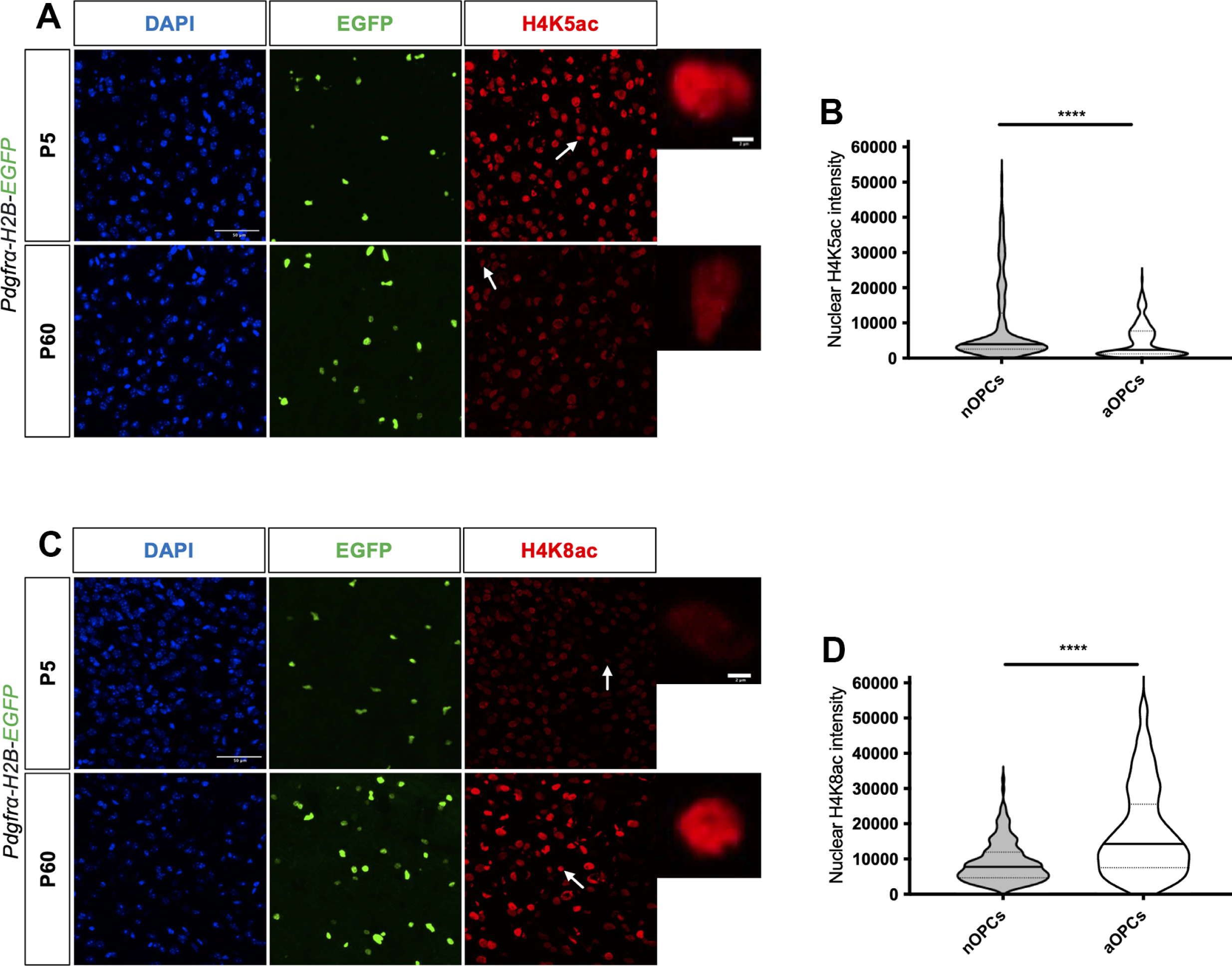
Differential levels of H4K5ac and H4K8ac intensity in the nuclei of OPCs in the developing and adult brain. **(A).** Immunohistochemical staining of the cortex of *Pdgfra-H2BEGFP* mice with H4K5ac specific antibody (red) in neonatal and adult brain sections. DAPI is stained in blue and PDGFRα^+^ cells are in green. Scale bar = 50 μm. **(B).** Violin plots of nuclear intensity of H4K5ac in PDGFRα^+^ cells in nOPCs and aOPCs. Data represent the distribution of the H4K5ac nuclear intensity measured in 200 cells from n = 3 mice (****p<0.0001, Student’s t test). **(C).** Immunohistochemical staining of the cortices of Pdgfra-H2BEGFP mice with H4K8ac specific antibody (red) in neonatal and adult brain sections. DAPI is stained in blue and PDGFRα^+^ cells are in green. Scale bar = 50 μm. Scale bar = 50 μm. **(D).** Violin plots of nuclear intensity of H4K8ac in PDGFRα^+^ cells in nOPCs and aOPCs. Data represent H4K8ac nuclei intensity of 200 cells from n = 3 mice (****p<0.0001, Student’s t test).

### Adult OPCs are characterized by increased H4K8ac chromatin occupancy at specific loci related to oligodendrocyte-specific genes

To begin addressing the functional relevance of specific histone marks in aOPCs, we interrogated the genomic distribution of the H4K8ac mark for loci with differential occupancy in the nuclei of nOPCs and aOPCs, using ChIP-sequencing analysis **(Fig. 5A)**. We used ChIP-grade antibodies specific for H4K8ac, that were initially validated using western blot (**Supplemental Fig. S3A**), and antibody arrays (**Supplemental Fig. S3B**), according to the ENCODE guidelines and obtained high quality reads from the six samples (three independent biological chromatin samples from aOPCs and three from nOPCs, **Supplemental Fig. S3C**). A principal component analysis of the data revealed a clear separation of the peaks from aOPC and nOPC samples **(Supplemental Fig. S3D)**. When the distribution was plotted relative to the transcriptional start site, we detected a similar pattern, but a much greater H4K8ac genomic occupancy in aOPCs than nOPCs **(Fig. 5B**), nicely correlating with the higher levels of H4K8ac detected in aOPCs. We next performed differential binding analysis to identify genomic locations differentially bound by H4K8ac between these two cell populations and identified 43,888 ChIP peaks with significantly greater occupancy in aOPCs **(Fig. 5C).** When the peak distribution was analyzed using the genomic features, we noted that 43.2 % of the peaks localized at the promoter region, 34.2 % at introns and 18.3 % at distal intergenic regions **(Fig. 5D).** To assign biological meaning to the chromatin regions with greater deposition of H4K8ac marks in aOPCs, we used Genomic Regions Enrichment of Annotations Tool (GREAT) (McLean et al., 2010). This analysis revealed biological processes related to transcription, protein folding and mitochondrial respiratory chain complex among the 9,491 genes with higher levels of the H4K8ac mark at the promoter region and biological processes related to axonal ensheathment and myelination for the remaining genes with differentially acetylated peaks localized intronic (2,442 genes) and distal intergenic (1,486 genes) regions **(Fig. 5E).** Together, these results identify H4K8ac occupancy in aOPCs at chromatin loci regulating transcription and myelination.

**Figure 5.**
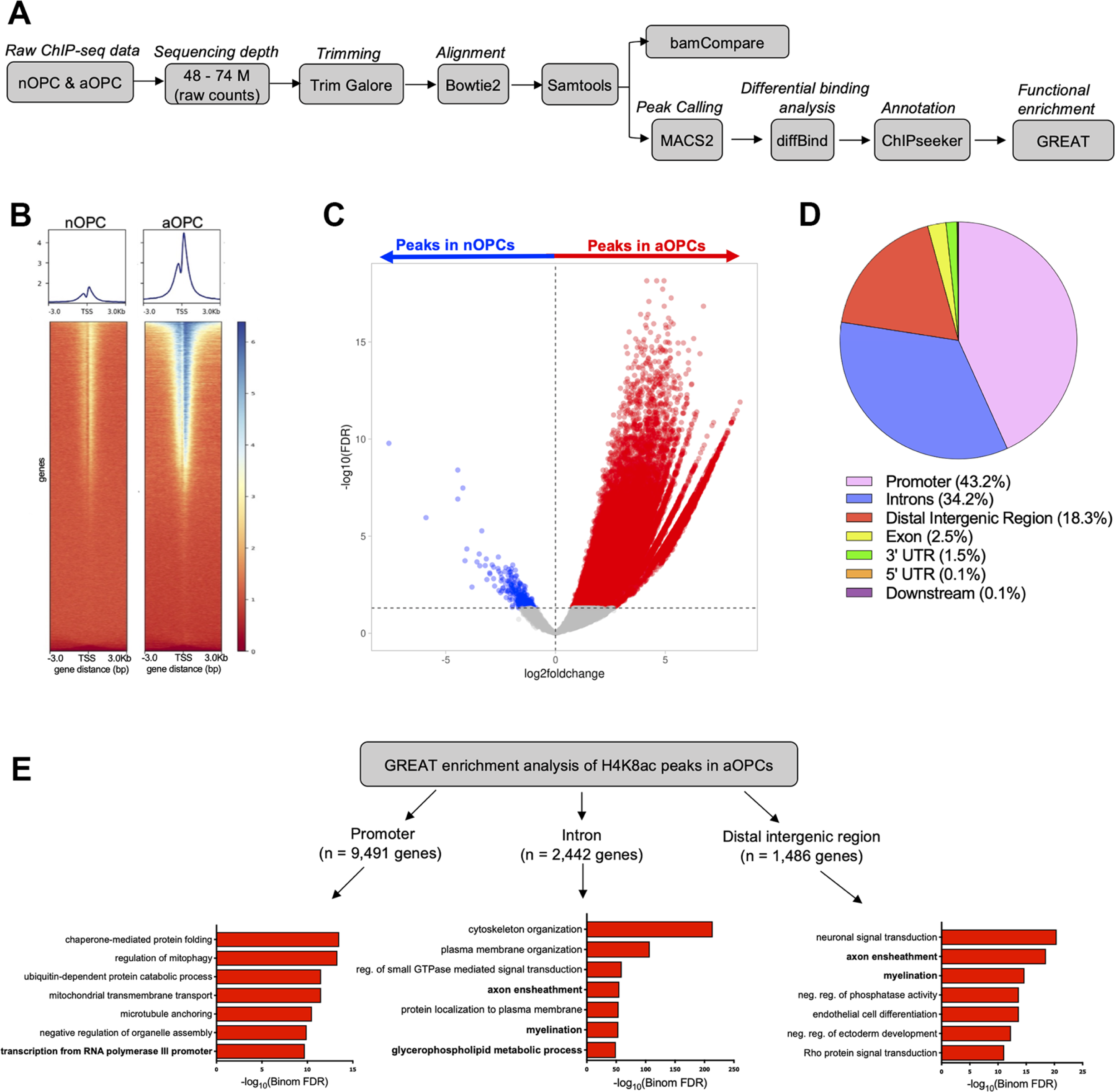
Adult oligodendrocyte progenitors are characterized by increased H4K8ac chromatin occupancy. **(A).** Experimental flowchart for the H4K8ac ChIP-sequencing data. **(B).** Genomic heatmaps showing the distribution of H4K8ac peaks in nOPCs and aOPCs at 3 kilobases (kb) around the transcription start site (TSS). **(C).** Volcano plot showing the differentially called H4K8ac peaks in aOPCs vs nOPCs (significance cut-off at FDR < 0.05). Differentially called peaks with greater occupancy in samples from aOPCs are shown in red while those with greater occupancy in nOPCs are shown in blue. **(D).** Pie chart showing the genomic distribution of the H4K8ac differential peaks present in aOPCs. **(E).** Functional annotation of H4K8ac differentially peaks in aOPCs at promoter, intron, and distal intergenic regions.

To better understand the transcriptional consequences of the greater genomic occupancy of the histone H4 mark in aOPCs, we overlapped the genes with greater H4K8ac genomic occupancy (FDR < 0.05, log_2_FC [aOPC/nOPC] >0.75). We noted that over 80% of the transcripts with higher levels in aOPCs corresponded to genes with chromatin regions bearing the H4K8ac mark **(Fig 6A)**. Ontology analysis of these genes identified categories related to regulation of transcription from RNA polymerase II (30.5%), myelin/lipid metabolic process (16.5%), and protein transport (13.6%), among the most prominent categories **(Fig. 6B).** Visualization of genomic occupancy led to highly reproducible patterns of peak enrichment in aOPCs (compared to nOPCs) in genomic regions corresponding to *Cnp* **(Fig. 6C),** *Mog* **(Fig. 6D),** and also other myelin proteins, such as *Plp1* **(Fig. 6E)** and *Mbp* **(Fig. 6F)**, the transcription factor *Nkx6-2* **(Fig. 6G),** and the lipid enzyme *Ptgds* **(Fig. 6H)**.

**Figure 6.**
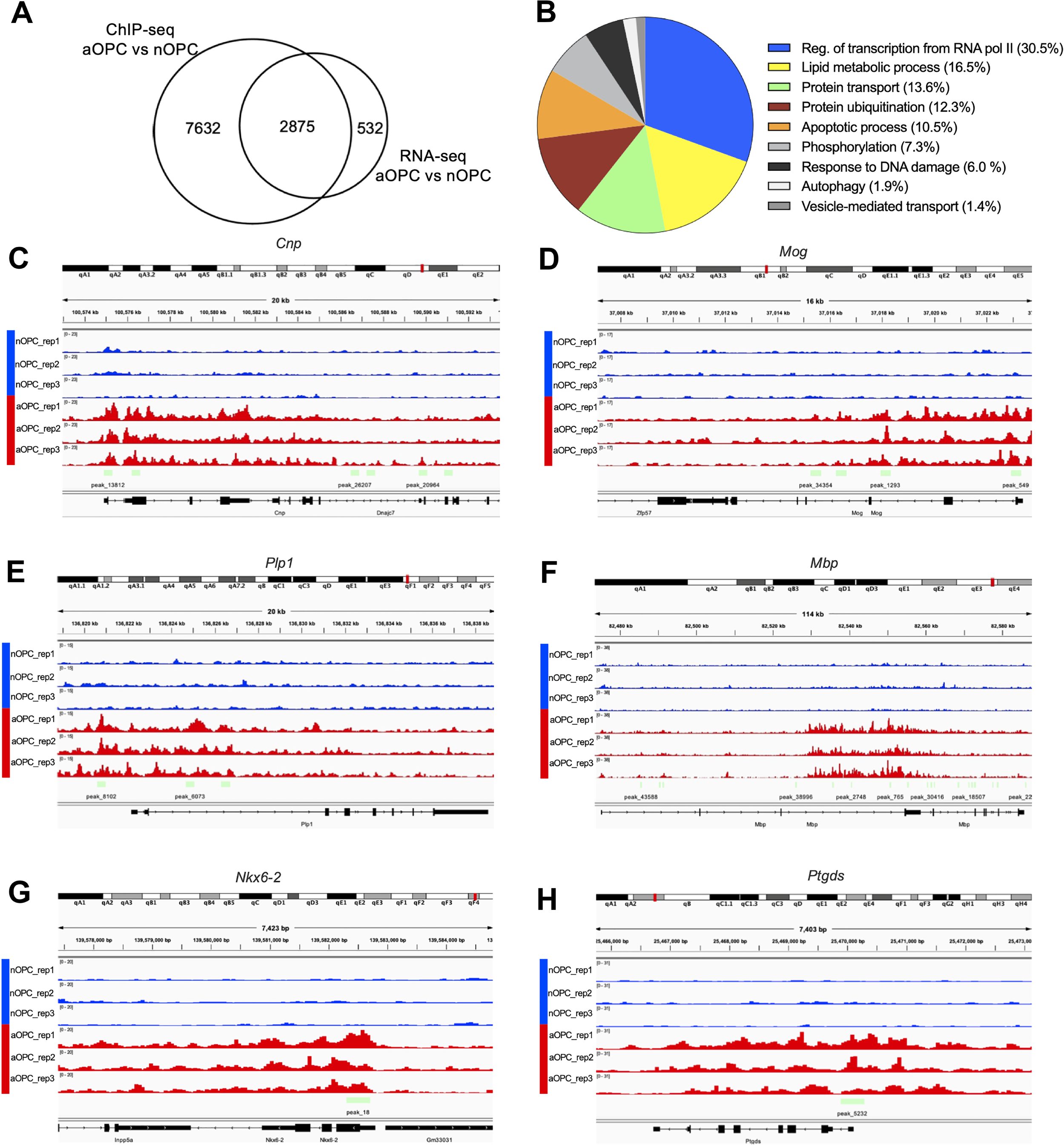
H4K8ac occupies genomic regions of genes expressed in mature oligodendrocytes in aOPCs. **(A).** Venn diagram showing the number of genes that overlapped between the H4K8ac ChIP-seq dataset and the RNA-seq data from aOPCs compared to nOPCs (FDR < 0.05, log_2_FC [aOPC/nOPC] >0.75). **(B).** Pie-chart showing the proportions of unique genes from the overlap, involved in each of the indicated biological processes. **(C-H).** Screenshots of Integrative Genomics Viewer (IGV) tracks (Robinson et al., 2011) showing H4K8ac enrichment in aOPCs (red) compared to nOPCs (blue) at the indicated loci, including *Cnp* **(C),** *Mog* **(D),** *Plp1* **(E),** *Mbp* **(F)**, *Nkx6.2* **(G)** and *Ptgds* **(H).** For each panel, the RefSeq gene track is shown at the bottom in black.

Together, these data are consistent with the interpretation of H4K8ac as an important activating histone mark, occupying chromatin regions corresponding to genes highly expressed in aOPCs.

### Blocking H4K8ac mark deposition with pharmacological inhibitors in aOPCs prevents the expression of genes related to the oligodendrocyte state

To functionally validate the transcriptional consequences of H4K8ac occupancy in aOPCs, we used pharmacological inhibitors of specific histone acetyltransferases responsible for H4K8ac deposition, such as KAT2B and KAT5 (Kim et al., 2020; Kimura & Horikoshi, 1998), which are expressed in the oligodendrocyte lineage and whose levels are higher in aOPCs than nOPCs (**Supplemental Fig. S4**). Garcinol – an inhibitor specific for KAT2B and KAT3B (Balasubramanyam et al., 2004) and NU-9056 - specific for KAT5, KAT2B and KAT3B (Coffey et al., 2012) were used in combination in order to decrease the levels of H4K8ac in aOPCs. H4K8ac nuclear intensity was used to validate the effectiveness of the treatment in reducing this histone mark in aOPCs **(Fig. 7A-B).** Functionally, the pharmacological inhibition of H4K8ac deposition did not impact the levels of the cell cycle related *Ccnd2* **(Fig. 7C),** but led to decreased levels of several transcripts characteristic of the differentiated state, such as *Nkx6-2* **(Fig. 7D)***, Ptgds* **(Fig. 7E)**, as well as differentiation genes such as *Cnp* **(Fig. 7F)**, while late myelin gene transcript levels were not affected **(Fig. 7G-I).** Together, these results suggest that the H4K8ac histone mark in aOPCs is responsible for the regulation of some genes characteristically found in mature oligodendrocytes, while also highlighting the fact that other mechanisms are likely at play to prevent these cells from differentiating and maintaining a proliferative state, albeit reduced compared to nOPCs.

**Figure 7.**
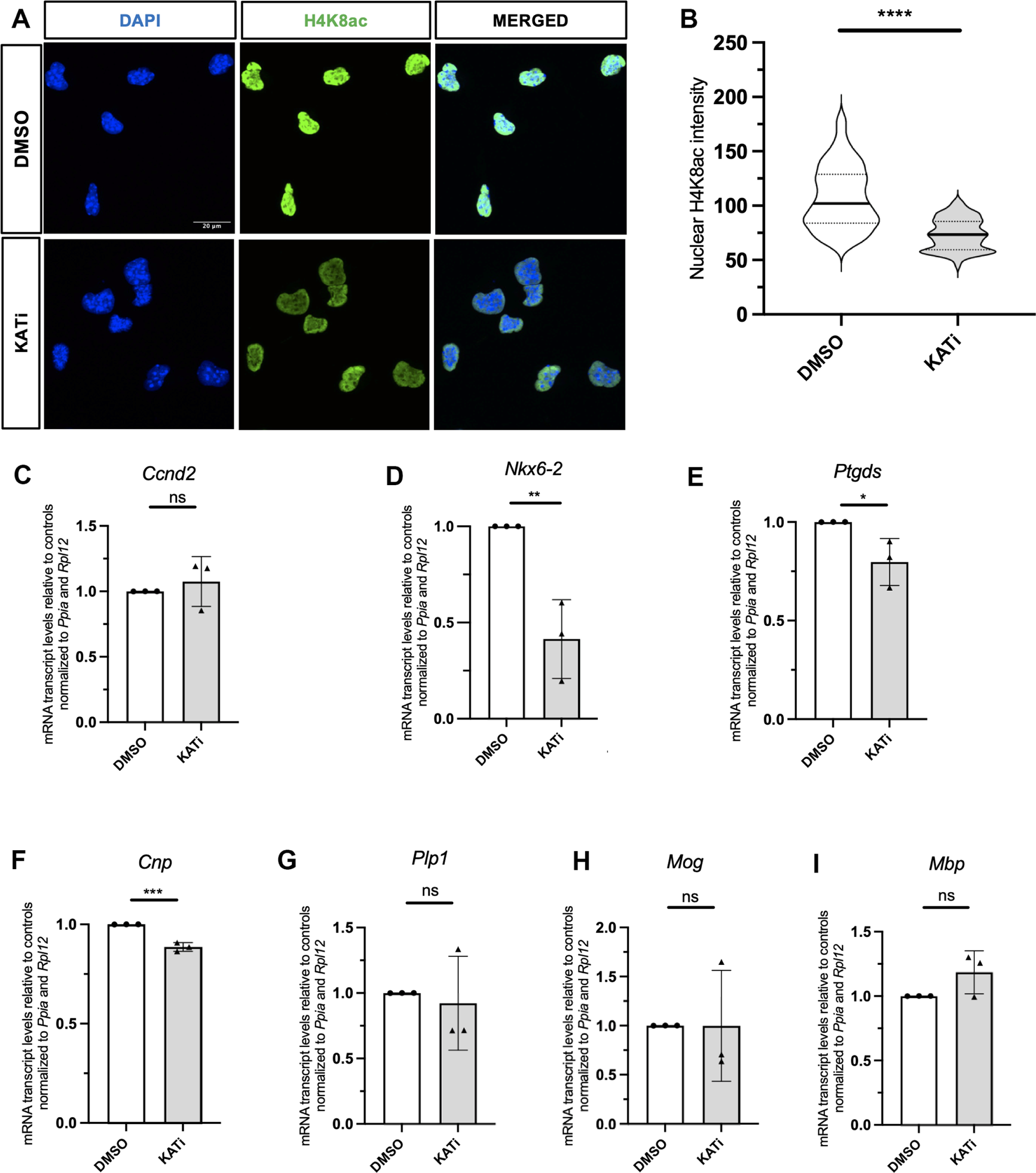
Pharmacological inhibition of H4K8ac deposition in aOPCs results in the downregulation of the transcript levels of genes characteristically expressed in mature oligodendrocytes. **(A)**. Representative confocal images of aOPCs treated either with DMSO or KATi (5 μM Garcinol and 0.2 μM NU-9056) for 48 h and stained with DAPI (blue) and H4K8ac (green). Scale bar = 20 μm. **(B).** Violin plots showing nuclear intensity of H4K8ac in KATi treated aOPCs (gray) and controls (white). Data represent H4K8ac nuclei intensity of 100 cells, n = 3 (****p<0.0001, Student’s t test). **(C-I).** The bar graph indicate the average transcript levels of Ccnd2 (C), Nkx6-2 (D), Ptgds (E), Cnp (F), Plp1 (G), Mog (H) and Mbp (I) in KATi treated aOPCs (gray bars) relative to controls (white bars). Data represent average values normalized to the geomean of the indicated reference genes and presented as relative to the levels in nOPC samples ± SD for n = 3 (*p < 0.05, **p<0.01, ***p<0.001, Student’s t test).

### Pharmacological inhibition of the H4K20me3 mark deposition in aOPCs promotes the expression of genes related to the mature oligodendrocyte state

As also the heterochromatic repressive mark H4K20me3 was higher in aOPCs than nOPCs, we asked whether the expression of oligodendrocyte differentiation genes could be regulated by the deposition of this mark in aOPCs by the enzymes SUV420H1 and SUV420H2. Using A-196 (SUV420i), a potent and selective pharmacological inhibitor of the lysine methyltransferases SUV420H1 and SUV420H2 (Bromberg et al., 2017), we monitored decreased H4K20me3 as metric of effectiveness of the pharmacological inhibition in treated aOPCs **(Fig. 8A-B).** The decreased H4K20me3 levels functionally resulted in decreased transcript levels of the cell cycle gene, *Ccnd2* **(Fig. 8C),** unchanged levels of *Nkx6.2* **(Fig. 8D),** *Cnp* **(Fig. 8E),** and *Mog* **(Fig. 8F),** and increased transcript levels of mature oligodendrocyte-specific genes *Plp,1 Mbp* and *Ptgds* **(Fig. 8G-I).** Taken together, these results suggest that the repressive H4K20me3 halts the expression of some myelin genes and contributes to the maintenance of the progenitor state in aOPCs.

**Figure 8.**
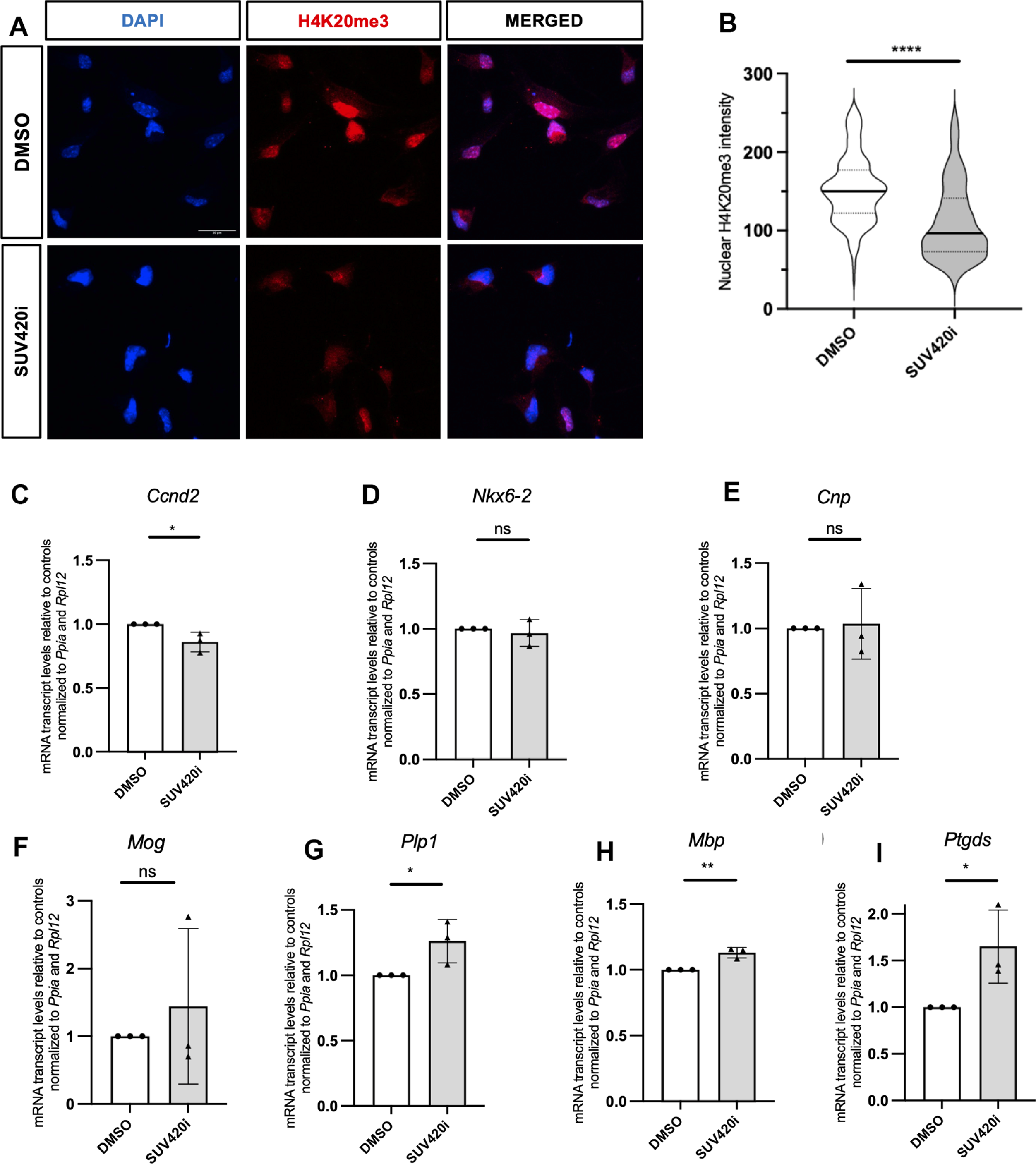
Inhibition of H4K20me3 deposition in aOPCs results in decreased levels of cell proliferation transcripts and increased levels of oligodendrocyte-specific genes. **(A)**. Representative confocal images of aOPCs treated either with DMSO or SUV420i (5 μM A-196) for 48 h and stained with DAPI (blue) and H4K20me3 (red). Scale bar = 20 μm. **(B).** Violin plots representing the distribution of nuclear intensity of H4K20me3 in SUV420i inhibitor treated aOPCs (gray) and DMSO controls (white). Data represent H4K20me3 nuclei intensity of 200 cells, n = 3 (****p<0.0001, Student’s t test). **(C-H).** The bar graphs indicate Ccnd2 (C), Nkx6-2 (D), Cnp (E), Mog (F), Plp1 (G), Mbp (H) and Ptgds (I) transcript levels in SUV420i treated aOPCs relative to controls. Data represent average values normalized to the geomean of the indicated reference genes and presented as relative to the levels in nOPC samples ± SD for n = 3 (*p < 0.05, **p<0.01, Student’s t test).

## DISCUSSION

While aOPCs represent the most abundant progenitor population of the adult brain, still relatively little is known about the molecular mechanisms that make these cells distinct from nOPCs. It is well accepted that these two cell populations share immunoreactivity for progenitor markers and differ from them for their ability to proliferate, migrate and respond to external signals (Chari et al., 2003; Gensert & Goldman, 2001; G. Lin et al., 2009; Wolswijk et al., 1990; Wolswijk & Noble, 1989). Although the molecular mechanisms underlying these functional differences are not completely understood, several studies identified important transcriptional and proteomic differences between these two states (de la Fuente et al., 2020; Moyon et al., 2021; Neumann et al., 2019; Spitzer et al., 2019). Using oligodendrocyte progenitor reporter mice to directly sort cells from neonatal and adult brains, we first validated transcriptional and functional differences between the two states. Functionally, we noted lower cell proliferation in aOPCs compared to nOPCs, which was consistent with the downregulation of genes related to cell cycle and proliferation. In addition, we detected several upregulated transcripts typically related to the mature oligodendrocyte state, including the transcription factor *Nkx6.2*, previously reported to modulate the expression of paranodal genes (Southwood et al., 2004), and myelin proteins, such as *Cnp, Mog, Plp1* and enzymes like *Aspa* and *Ptgds*, which were also identified in previous studies (de la Fuente et al., 2020).

As the epigenetic landscape is unique to each cell type and developmental stage, we sought to identify the histone marks with differential abundance in aOPCs compared to nOPCs and adopted an unbiased proteomic analysis to identify histone post-translational modifications (PTMs) in these two cell types. Of the several differences in histone H4 post-translational modifications, we further validated lower levels of H4K5ac in aOPCs, a mark which we previously reported to decrease during the early stages of nOPC differentiation and for cross talk with arginine related marks, such as H4R3me2s (Scaglione et al., 2018). We also detected and validated higher levels of H4K8ac, a mark previously associated with active transcription initiation and elongation (Agalioti et al., 2002; Radzisheuskaya et al., 2021; Wang et al., 2008).. We further identified high levels of the repressive heterochromatic H4K20me3 mark (Gabellini & Pedrotti, 2022; Schotta et al., 2004) in aOPCs.

The genome-wide distribution of H4K8ac in aOPCs was predominantly observed at promoter regions of genes with higher transcript levels in these cells, such as *Nkx6-2* transcription factor and lipid metabolic enzyme *Ptgds*. Greater H4K8ac occupancy was also found at introns and distal intergenic regions of myelin genes such as *Cnp, Plp, Mog* and *Mbp,* a distribution which was consistent with previous studies identifying this mark at enhancer regions (Goudarzi et al., 2016; Q.-L. Li et al., 2019). The detection of greater H4K8ac occupancy in aOPCs occurring at loci corresponding to oligodendrocyte stage-specific genes was also consistent with previous studies reporting the transcriptional signature of aOPCs to be more similar to mature oligodendrocytes than to nOPCs (Moyon et al., 2015) and with proteomic studies identifying oligodendrocyte specific proteins in aOPCs (de la Fuente et al., 2020). The strong correlation between transcript levels in aOPCs and chromatin regions with high H4K8ac occupancy supported the transcriptional activation role previously reported for this mark (Gupta et al., 2017; Q.-L. Li et al., 2019). It was further supported by the fact that pharmacological inhibition of the enzymes responsible for the deposition of H4K8ac lowered the transcript levels of the gene targets in vitro. However, the precise mechanism of activation in aOPCs is not entirely clear. Previous studies had suggested that transcriptional activation by histone acetylation was mediated by the recruitment of the transcription initiation complex by the bromodomain protein TAFIID (Jacobson et al., 2000) and the facilitation of the elongation complex by BRD4 (Muhar et al., 2018), which requires dual acetylated sites on histone H4K5ac and K8ac to be recruited (Filippakopoulos et al., 2012; Zaware & Zhou, 2019). While we noted that H4K5ac is less abundant in aOPCs compared to nOPCs, it is conceivable that the presence of the two marks may account for the recruitment of BRD4 to specific genomic region encoding for the genes, whose transcripts are upregulated in aOPCs.

It is important to note here that, as previously reported, besides myelin genes, aOPCs are also characterized by genes characteristic of the mature oligodendrocyte state (such as *Aspa* and *Car2*) and yet those cells are less efficient in generating myelinating oligodendrocytes than nOPCs. The identification of the co-occurrence of high levels of the repressive mark H4K20me3 (Karachentsev et al., 2005) in aOPCs, may provide a potential explanation. Pharmacological inhibition of the enzymes responsible for H4K20me3 deposition resulted in increased transcript levels of genes expressed in mature oligodendrocytes, a finding consistent with the idea that in aOPCs the repressive H4K20me3 mark may serve the function of halting differentiation while repressing a complete exit from the cell cycle. Future work may test the hypothesis that H4K20me3 mark may keep aOPCs at a progenitor state while also creating heterochromatic regions which may render the process of differentiation towards myelinating OL, harder to achieve. It would be interesting to determine whether down-regulating the levels of H4K20me3 in aOPCs after demyelination, possibly in association with additional epigenetic drugs, could improve remyelination.

Of interest is the consideration that the repressive H4K20me3 histone mark has also been previously related to senescence, as its levels increase in aging (Sarg et al., 2002) as well as in senescent (Nelson et al., 2016) and progeroid cells (Shumaker et al., 2006).

Together, these data identify histone H4 post-translational modifications, such as H4K8ac and H4K20me3 as important for the definition of the transcriptional features and functional properties of aOPCs, although future studies will be needed to carefully define all the other modifications that render the epigenetic landscape of these cells unique.

## MATERIALS AND METHODS

### Mouse models

All experiments were performed according to the Institutional Animal Care and Use Committee (IACUC)-approved protocols of the Advanced Science Research Center (ASRC), City University of New York (CUNY). Mice were maintained in a temperature- and humidity-controlled facility on a 12-h light–dark cycle with food and water *ad libitum*. Mice from either sex were used for all experiments*. Pdgfrα-H2B-EGFP* reporter mouse line was used to sort for oligodendrocyte progenitor cells, and for immunohistochemistry in the neonatal (P2-P5) and adult (P60) brains (Klinghoffer et al., 2002).

### Fluorescence-activated cell sorting (FACS)

Oligodendrocyte progenitors were isolated from the forebrains of postnatal day 5, and 60 *Pdgfrα-H2BEGFP* mice using fluorescence-activated cell sorting (Moyon et al., 2021). The tissues were dissected in HBSS 1X (HBSS 10X (Invitrogen), 0.01 M HEPES buffer, 0.75% sodium bicarbonate (Invitrogen), and 1% penicillin/streptomycin followed by mechanical and enzymatic dissociation (30 μg/ml of papain in DMEM-Glutamax, with 0.24 μg/ml L-cysteine and 40 μg/ml DNase I) to obtain cell suspension. The cell preparation was then carefully layered on top of a pre-formed Percoll density gradient and centrifuged for 15 min at 20,000 *g* at 4 °C. Cells were collected and stained with propidium iodide (PI) for 2 min at RT. GFP-positive and PI-negative cells were sorted by fluorescence-activated cell sorting (FACSAria^TM^, Becton Dickinson Biosciences). For OPC isolation in the adult mice, only the population with strongest GFP+ signal was collected as true aOPCs.

### Fluorescence-activated nuclei sorting (FANS)

Nuclei of oligodendrocyte progenitors were sorted from frozen forebrains harvested from *Pdgfrα-H2B-EGFP* mice using fluorescence-activated nuclei sorting (Jiang et al., 2008). Frozen forebrain tissues were cut into small pieces and homogenized in lysis buffer (0.32 M sucrose, 3 mM Mg(Ac)_2_, 0.1 mM EDTA, 10 mM Tris HCL pH 8.0, 0.1 % TritonX-100) using a Dounce tissue grinder on ice. The homogenate was filtered with a 70 µm strainer and transferred into an ultracentrifuge tube (Beckman Coulter, 331372). Sucrose solution (1.8 M sucrose, 3 mM Mg(Ac)_2_, 10 mM Tris HCL pH 8) was added to the bottom of the tube drop by drop to form a gradient. The sample was centrifuged at 24,000 rpm for 1 h,15 min at 4 °C. The supernatant was discarded, the pellet resuspended in FACS buffer (2% FBS, 2mM EDTA, 0.2% Saponin, 0.01% Sodium azide in PBS) and then filtered with a 40 µm strainer. Nuclei were incubated with 4′,6-diamidino-2-phenylindole (DAPI) for 15 min on ice. GFP-positive and DAPI-positive nuclei were sorted by fluorescence-activated nuclei sorting. For aOPC nuclei isolation, only the population with strongest GFP+ signal was collected as true aOPCs.

### Primary oligodendrocyte progenitors cell isolation and culture

Primary cortical neonatal and adult OPCs were isolated from postnatal day 5 and day 60 C57BL/6 mice (purchased from Jackson Laboratories), following the protocols described in the Adult Brain Dissociation Kit (ABDK, Miltenyi, 130-107-677). Briefly, neonatal and adult mice were euthanized, the cortices dissected and immediately placed in cold Hibernate A Low Fluorescence (HALF) medium (Fisher Scientific, NC0285514). The cortices were cut into 0.5 cm pieces and placed in C-tubes (Miltenyi, 130-093-237) containing enzyme and buffers provided in the kit. The C-tubes containing the cortices were placed in the gentleMACS™ Octo Dissociator with Heaters for dissociation (Miltenyi, 130-094-427; Program - 37C_ABDK_01). The resulting cell suspension was strained using a 70 μm MACS SmartStrainer (Miltenyi, 130-098-462) and the filtrate centrifuged at 300×g for 10 min at 4°C. The supernatant was completely aspirated, and debris removal performed using the Debris Removal Solution (Miltenyi, 130-109-398). Next, red blood cells were lysed by resuspending cell pellet in cold Red Blood Cell Lysis Solution (Miltenyi, 130-094-183). To prevent non-specific binding to the antibodies, the cells were pre-incubated with the FcR Blocking Reagent (Miltenyi, 130-059-901) and then labeled with O4 antibody-conjugated microbeads (Miltenyi, 130-094-543). O4+ oligodendrocyte progenitors were isolated by applying the cell suspension onto an MS column (Miltenyi, 130-042-201) placed on a magnetic field separator. The magnetically labeled cells were washed with PBS/BSA buffer (Miltenyi, 120-091-376) three times and immediately flushed out with 1 mL of SATO media by firmly pushing the plunger into the column. The cells were cultured on poly-D-lysine (PDL) coated plates. The isolated primary OPCs were cultured in SATO medium (Dulbecco’s modified Eagle’s medium (DMEM), 100 μg/mL BSA, 100 μg/mL apo-transferrin, 16 μg/mL putrescine, 62.5 ng/mL progesterone, 40 ng/mL selenium, 10 μg/mL insulin, 1 mM sodium pyruvate, 5 μg/ml N-acetyl-cysteine, 10 ng/ml biotin, 5 μM forskolin, 1% GlutaMAX^TM^ Supplement (Gibco), B27 Supplement and Trace Element B) supplemented with Platelet-Derived Growth Factor-AA (PDGFA) (30 ng/ml) and basic Fibroblast Growth Factor (bFGF, Peprotech, 100-18B) (30 ng/ml) for the first day. Media change was performed on the second and all other days with SATO medium supplemented (PDGFA) (20 ng/ml) and basic Fibroblast Growth Factor (bFGF) (20 ng/ml).

### RNA extraction

For RNA-sequencing, RNA was isolated from washed FAC-sorted cells using an AllPrep DNA/RNA Micro Kit (Qiagen). RNA was extracted from cortical tissues from three biological replicates for each age using TRIzol (Invitrogen). For RT-qPCR, RNA was extracted from primary nOPCs and aOPCs using TRIzol (Invitrogen) and the RNeasy Mini Kit (Qiagen, 74106). RNA samples were resuspended in water and further purified with RNeasy columns with on-column DNase treatment (Qiagen). RNA purity was assessed by measuring the A260/A280 ratio with a NanoDrop, and RNA quality checked using an Agilent 2100 Bioanalyzer (Agilent Technologies).

### RNA-sequencing

For nOPCs, 85 ng of total RNA per sample was used for library construction with the Ultra-Low-RNA-seq RNA Sample Prep Kit (Illumina) and pair-end sequenced using the Illumina HiSeq 2000 instrument, according to the manufacturer’s instructions for paired end read runs ((Moyon et al., 2016). For aOPCs, 250 ng of total RNA per sample was used for library construction with the TruSeq RNA Sample Prep Kit (Illumina) and pair-end sequenced using the Illumina HiSeq 2500 instrument, according to the manufacturer’s instructions for paired end read runs (Moyon et al., 2021). High-quality reads were aligned to the mouse reference genome (mm10) using the subread package toolset (Liao et al., 2013). The featureCounts program (Liao et al., 2014) was used to count mapped reads and DESeq2 (Love et al., 2014) was used for differential gene expression analysis. We used a cut-off of FDR < 0.05 and |foldchange| > 0.75 to identify differentially expressed genes. For gene ontology analysis, we used DAVID ontology tool. The overall mapped reads and sample names are listed below. Data deposited in GEO, accession number GSE66047 for nOPCs and GSE122446 for aOPCs. *RNAScope In situ* hybridization was performed using RNAScope probe and detection reagents from Advanced Cell Diagnostics (ACD) in accordance with the manufacturer’s guidelines. Paraffin-embedded P5 (nOPCs) and P60 (aOPCs) wild-type mice brain sections were first de-paraffinized. RNAScope probes were hybridized for 2 h at 40 °C in a humidity-controlled oven (HybEZ II, ACDbio). The signal of target RNA was visualized through probe-specific horseradish-peroxidase-based detection. Slides were then counter-stained with DAPI, coverslipped with Prolong Gold Antifade (Thermofisher), and imaged with Zeiss LSM800 Confocal Microscope. The following RNAScope probes were used; *Pdgfra* (ACD, 480661-C3), *Olig2* (ACD, 447091), *Ptgds* (ACD, 492781-C2) and *Cnp* (472241).

### Quantitative real-time PCR

RNA was reverse transcribed with qScript cDNA Supermix (Quantabio, 95047). Quantitative real-time PCR (RT-qPCR) reactions were run with PerfeCTa SYBR GREEN FastMix, ROX reagent (Quantabio, 95,072) at the Epigenetics Core facility of ASRC. The expression levels for each transcript were normalized to the geometric mean of housekeeping genes: *Rpl12* and *Ppia.* Next, CT values of the technical replicates were average for every biological replicate and data presented as relative to the transcript levels in controls. The sequences of primers used for RT-qPCR are listed below:

*Ptgds:* F-TGCAGCCCAACTTTCAACAAG and R-TGGTCTCACACTGGTTTTTCCT *Nkx6-2:* F-AAGTCTGCCCCGTCTCAAC and R-GGTCTGCTCGAAAGTCTTCTC *Cnp:* F-TTCTGGAGATGAACCCAAGG and R-TGGGTGTCACAAAGAGAGCA *Plp1:* F-CCAGAATGTATGGTGTTCTCCC and R-GGCCCATGAGTTTAAGGACG *Ccnd2:* F-GAGTGGGAACTGGTAGTGTTG and R-CGCACAGAGCGATGAAGGT *Mog:* F-AGCTGCTTCCTCTCCCTTCTC and R-ACTAAAGCCCGGATGGGATA *Mag:* F-CTGCCGCTGTTTTGGATAATGA and R-CATCGGGGAAGTCGAAACGG *Mbp:* F-GGCGGTGACAGACTCCAAG and R-GAAGCTCGTCGGACTCTGAG *Ppia:* F-CACAAACGGTTCCCAGTTTT and R-TTCCCAAAGACCACATGCTT *Rpl12:* F-GAACAGACAGGCCCAGATCG and R-CAGACTGTGCAGTACCCAGG

### Chromatin isolation and chromatin immunoprecipitation

Fluorescence-activated nuclei sorting (FANS) was performed on frozen *Pdgfrα-H2B-EGFP* P5 and P60 forebrain homogenate after it was crosslinked with 1% paraformaldehyde and quenched with 125 mM glycine. The nuclear pellets were lysed with lysis buffer (50 mM Tris-HCl, 10 mM EDTA, 1% SDS, pH 8) and then sonicated with Bioruptor (Diagenode, 15 cycles, 30s ON/30s OFF). The size of the DNA in the sheared chromatin fragments was checked before precipitation by a Tapestation system (Agilent 4200) to ensure that the majority of fragment size was 200–400 bp. Protein A-Dynabeads (Thermo Fisher, 10002D) were washed three times with PBS/BSA buffer (5 mg/mL) and antibody-bead conjugation was carried out for 2 h on a rotator at 4°C. Immunoprecipitation was performed with 9 μg of anti-H4K8ac (Active Motif, 61103) per ChIP sample after preclearing chromatin with protein A-Dynabeads. As negative control, immunoprecipitation was performed with 15 ug of normal rabbit IgG (Sigma, NNI01-100UG). For each biological replicate, a total of 2.25 X 10^6^ nuclei were used for H4K8ac immunoprecipitation and 7.5 X 10^5^ nuclei were used for normal IgG immunoprecipitation. A total of three biological replicates were used per age. The immunoprecipitation reaction was performed overnight on a rotator at 4°C. The supernatants of the negative control samples were saved as Input while reverse crosslinking was performed on the H4K8ac immunoprecipitated chromatin using elution buffer (100 mM NaHCO_3_ & 1 % SDS) supplemented with proteinase K (Millipore Sigma, 3115887001, 20 mg/mL) and incubated at 68 °C in a thermomixer at 1300 rpm for 2 h. DNA extraction was then performed with Phenol:Chloroform:Isoamyl Alcohol (25:24:1) and precipitated with 3M Sodium acetate, 2.5% linear acrylamide carrier and cold 100% ethanol at −20 °C for 2 h. The precipitated DNA was washed with cold 70 % ethanol, dissolved in ddH_2_O and placed at 4 °C overnight for complete elution. DNA yield was then measured using Qubit 3 (High Sensitivity DNA Assay, ThermoFisher). Libraries were prepared using 2 ng of ChIP DNA and 10 ng of Input DNA (KAPA Hyper Prep Kit, KK8502). The input and ChIP samples were sequenced by Illumina HiSeq 4000. The filtered read counts of the samples are in **Supplemental Fig. 3**.

### ChIP-seq analysis

FASTQC was used for quality control. The adapter sequences were removed from the FASTQ files using Trim Galore (Krueger et al., 2021). Trimmed reads were aligned to the *Mus musculus* reference genome GRCm38 using Bowtie2 (v.2.4.5) program (Langmead & Salzberg, 2012). SAMTools (H. Li et al., 2009) was used to remove PCR duplicates and reads mapped to blacked list regions (Amemiya et al., 2019) and alignments with low mapping quality (q < 20). The deepTools bamCompare function (Ramírez et al., 2016) was used to normalize the ChIP samples to their respective input samples. The peaks in the ChIP sample in reference to the input sample in nOPCs and nOPCs were called from read alignments by MACS2 algorithm (Zhang et al., 2008). DiffBind (Stark & Brown, 2011) was used for the differential binding of H4K8ac in aOPCs compared to nOPCs. A total of 43,888 significantly differential sites were identified with FDR < 0.05. Annotation of the H4K8ac peaks was performed with ChIPseeker in R package (G. Yu et al., 2015) and the functional enrichment of annotation performed with GREAT tool (McLean et al., 2010). Finally, the coverage of normalized ChIP-Seq data were visualized with the integrative genomics viewer (IGV) program (Robinson et al., 2011).

### Histone extraction

Histones were extracted from fluorescence activated sorted OPC nuclei, using the acid extraction method (Shechter et al., 2007). A total of 5 X 10^6^ OPC nuclei per biological replicate was suspended into 400 µL of 0.4 N H_2_SO_4_ and incubated overnight at 4 °C on a rotator. After incubation, a 10 min centrifugation at 16,000 *g* 4 °C was used to remove nuclear debris. The supernatant containing the histones was transferred into a new tube and precipitated with 20 % trichloroacetic acid (TCA) for 1 h on ice. Histones were pelleted by centrifugation at 16,000 *g* for 10 min at 4 °C. The supernatant was discarded, and histone pellets washed twice with 1 mL of cold acidified acetone (0.1 % HCl) and then with 1 mL of cold acetone. The pellets were left to air-dry on ice for 1 h after which the pellets were resuspended in 100 µL of ddH_2_O by pipetting up and down and with mild vortexing. The histones were left to fully dissolve overnight at 4 °C.

### Quantitative histone proteomics analysis

Unbiased proteomics analysis was carried out on extracted histones from fluorescence-activated nuclei sorted oligodendrocyte progenitors. The extracted histones were first derivatized using propionic anhydride as described (Garcia et al., 2007). Briefly, the sample was diluted with methanol 1:1 (v:v) at pH 8.0. The propionic anhydride reagent mixture was prepared by mixing 75 µL propionic anhydride with 25 µL methanol. An aliquot equal to one quarter of the total volume of the histone sample was added and ammonium hydroxide was immediately used to titrate the pH back to 8.0 if necessary. The reaction was allowed to incubate for 20 min. The sample was dried to approximately 5 µL and in the Speedvac and the derivatization repeated as described. After another concentration using the Speedvac, the samples were resuspended in ammonium bicarbonate and digested overnight with 0.2 µg of sequencing grade modified trypsin (Promega) at 37 °C. The samples were again concentrated using the Speedvac and resuspended in 50 % methanol as above and the propionic anhydride derivatization repeated as described above. The dried sample was resuspended in 0.5 % acetic acid and the samples were loaded onto equilibrated microspin Harvard apparatus using a microcentrifuge and rinsed three times with 0.1% trifluoroacetic acid (TFA). The extracted samples were further washed with 0.5 % acetic acid and eluted with 40 % acetonitrile in 0.5% acetic acid followed by the addition of 80 % acetonitrile in 0.5 % acetic acid. The organic solvent was removed using a Speedvac, reconstituted in 0.5 % acetic acid, and then analyzed by liquid chromatography – mass spectrometry (LC-MS). LC separation was performed online on an EASY-nLC 1200 (Thermo Scientific) utilizing Acclaim PepMap 100 (75 µm x 2 cm) precolumn and PepMap RSLC C18 (2 µm, 100A x 50 cm) analytical column. Peptides were gradient eluted from the column directly into an Orbitrap Eclipse (Thermo Scientific) mass spectrometer using a 120 min acetonitrile gradient from 5% to 60% B followed by a ramp to 100% B in 20 min and final equilibration in 100% B for 15 min (A=2% ACN 0.5% AcOH / B=80% ACN 0.5% AcOH). Flowrate was set at 200 nL/min. High resolution full MS spectra were acquired with a resolution of 120,000, an AGC target of 4e5, with a maximum ion time of 50 ms, and scan range of 400 to 1500 m/z. All MS/MS spectra were collected using the following instrument parameters: resolution of 30,000, AGC target of 2e5, maximum ion time of 200 ms, one microscan, 2 m/z isolation window, fixed first mass of 150 m/z, and NCE of 27.

MS/MS spectra were searched against a Uniprot *Mus musculus* database using Sequest within P.D 1.4 for entire database analysis and Byonic for H3.1 and H4 PTM analysis. The search parameters for Sequest were as follows: mass accuracy better than 10 ppm for MS1 and 0.02 Da for MS2, two missed cleavages, fixed modification carbamidomethyl on cysteine, variable modification of oxidation on methionine. The data was filtered using a 1 % FDR cut off for peptides and proteins against a decoy database and only proteins with at least 2 unique peptides were reported. For the PTM analysis on histone H3.1 and H4 the search parameter was as follows: mass accuracy better than 10 ppm for MS1 and 0.02 Da for MS2, two missed cleavages, fixed modification carbamidomethyl on cysteine, variable modification of oxidation on methionine, monomethylation and propionic anhydride, dimethylation, trimethylation and acetylation on lysine residues, mono and demethylation on arginine, and phosphorylation on serine, threonine and tyrosine residue. For data analysis, the area under the curve for each peptide isoform was integrated and all the areas for a given peptide isoform were summed and represented as 100 % of the peptide intensity for a given peptide sequence. The percentage area allocated for each peptide isoform was calculated and the average percentage calculated for nOPCs and aOPCs. The fold change of the average abundance of aOPCs vs nOPCs was then calculated and those modifications which showed consistency across samples, and which were represented in more than two peptide sequences and showing the same behavior, were further analyzed.

### Western blot analysis

Western blotting was carried out on histone extracts from FAC-sorted nOPCs and aOPCs by sodium dodecyl sulfate–polyacrylamide gel electrophoresis (SDS-PAGE) as previously described (Dansu et al., 2022). A wet transfer of the proteins onto a 0.2 μm polyvinylidene fluoride (PVDF) membrane (Biorad, 1620177) was then carried out. The membranes were blocked for 1 h in 5 % non-fat dry milk (NFDM), 0.1 % Tween in Tris-buffered saline (TBS). Primary antibodies were incubated overnight at 4°C in 5 % milk, 0.1 % Tween/TBS. Membranes were washed with 0.1% Tween/TBS and incubated at RT for 1 h with horseradish peroxidase conjugated secondary antibodies (Jackson Immunoresearch, 1:10,000) in 5 % milk, 0.1% Tween/TBS. ECL Prime Western Blotting Detection Reagent kit (GE Healthcare, RPN2232) were then used to develop the membrane. ImageJ was used to quantify the intensity of the histone bands. The antibodies used for western blotting are: H3K9ac (Abcam, ab4441, 1:2000), H3K18ac (Abcam, ab1191,1:2000), H3K27ac (Abcam, ab4729, 1:2000), H3K4me3 (Abcam, ab8580, 1:2000), H3K9me3 (Abcam, ab8898, 1:1000), H3K27me2 (Cell Signaling, 9728s, 1:2000), H3K27me3 (Diagenode, C1541095, 1:2000), Total H3 (Cell Signaling, 12648S, 1:5000), H4K5ac (Active Motif, 39699, 1:2000), H4K8ac (Active Motif, 61103, 1:2000), H4K12ac (Active Motif, 39927, 1:2000), H4K16ac (Abcam, ab109463, 1:2000), H4K20me3 (Millipore, 07-463, 1:2000), Histone H4 (Abcam, ab197517, 1:5000).

### Immunohistochemistry (IHC) and Immunocytochemistry (ICC)

Immunohistochemistry was performed as previously described(Marechal et al., 2022). Mice were perfused with 4% paraformaldehyde (PFA), brains dissected and left immersed in fixative (4 % PFA) overnight at 4°C. The fixed brains were cryo-protected in sucrose solutions, and frozen embedded in OCT (Optimum Cutting Temperature, Tissue-Tek). Tissue sections (12 μm) were permeabilized and blocked by incubation with PGBA (0.1 M pH 7.4, 0.1% gelatin porcine type A, 1% BSA, and 0.002% Sodium azide), 10% normal goat or donkey serum and 0.1% Triton X-100 at room temperature (RT) for 1 h. Sections were incubated overnight at 4°C with primary antibodies, rinsed and then incubated with Alexa-Fluor™ secondary antibodies for 1 h at RT. Stained tissues were cover-slipped in DAPI Fluoromount G mounting medium (SouthernBiotech, 0100-20), allowed to dry, and imaged with Zeiss LSM800 Confocal Microscope. ImageJ was used to quantify immunofluorescence intensity. For immunocytochemistry, cells were fixed with 4 % PFA for 20 min at RT and non-specific binding blocked by incubation in PGBA, 10% normal goat serum, and 0.1% Triton X-100 1 h at RT. The fixed cells were then incubated with primary antibodies overnight at 4 °C, rinsed and then incubated with Alexa-Fluor™ secondary antibodies for 1 h at RT. Stained cells were cover-slipped in DAPI Fluoromount G mounting medium (SouthernBiotech, 0100-20), allowed to dry, and imaged with Zeiss LSM800 Confocal Microscope. ImageJ was used to quantify immunofluorescence intensity. The antibodies used for immunohistochemistry and immunocytochemistry and relative dilutions as follows: OLIG2 (Millipore, MABN50, 1:500), OLIG2 (Millipore, AB9610, 1:500), H4K5ac (Active Motif, 39699, 1:1000), H4K8ac (Abcam, ab15823, 1:1000), H4K20me3 (Millipore, 07-463, 1:500), 488-anti-rabbit (Invitrogen, A11034, 1:500), 488-anti-mouse (Invitrogen, A28175, 1:500), 555-anti-rabbit (Invitrogen, A21428, 1:500) and 555-anti-mouse (Invitrogen, A21422, 1:500).

### EdU incorporation assay

Click-iT™ EdU Cell Proliferation Kit (Invitrogen, C10337) was used to measure cell proliferation as descried in the protocol. Briefly, isolated primary nOPCs and aOPCs were plated in PDL-coated 8-well chamber slides and allowed to recover overnight (Thermo Fisher, 154739). The cells were incubated with 10 µM EdU for 5 h. The cells were then washed and fixed with 4% PFA for 15 min at RT, washed twice with 3% BSA in PBS and permeabilized with 0.5% Triton for 20 min at RT. The permeabilization buffer was removed and cells washed twice with 3 % BSA in PBS. Next, 0.5 mL of Click-iT® reagent was added and incubated in the dark for 30 min at RT. The cells were washed once with 3 % BSA in PBS and wells removed. The slide was then cover-slipped in DAPI Fluoromount G mounting medium (SouthernBiotech, 0100-20), allowed to dry, and imaged with Zeiss LSM800 Confocal Microscope.

### Pharmacological inhibition of KATs and SUV420H1/H2 in aOPCs

For KAT inhibition experiment, primary aOPCs were treated with a combination of KAT inhibitors (Garcinol, 5 μM and NU-9056, 0.2 μM) for 48 h. For SUV420H1/H2 inhibition, primary aOPCs were treated with the highly specific SUV420H1/H2 specific inhibitor (A-196, 5 μM) for 48 h. Control cultures were treated with dimethyl sulfoxide (DMSO). The dose and duration of the inhibitors used were selected based on previous studies conducted cells (Bromberg et al., 2017; Scaglione et al., 2018) and/or after initial dose-response cytotoxicity screen. Garcinol was purchased from Abcam (ab141503), NU-9056 from Sigma (5005110001) and A-196 from Sigma Millipore (SML1565).

### Gene Ontology analysis

Gene ontology (GO) analysis of the genes differentially expressed in aOPCs compared to nOPCs was performed with DAVID GO ontology tool (Huang et al., 2009; Sherman et al., 2022). Functional analysis of the differentially called H4K8ac peaks in aOPCs compared to nOPCs was carried out with GREAT tool (McLean et al., 2010).

### Quantification and statistical analysis

Confocal images were analyzed using Fiji-ImageJ. Volcano-plots were generated with VolcaNoseR (Goedhart & Luijsterburg, 2020). Integrative genomics viewer (IGV) program (Robinson et al., 2011) was used to visualize chromatin occupancy of H4K8ac ChIP-seq peaks on the genome. All other graphs were generated with GraphPad Prism and used for statistical analysis. Data are represented as mean ± standard deviation (SD).

## DATA AVAILABILITY

The mass spectrometry raw files related to histone proteomic data from nOPCs and aOPCs are accessible at www.massive.ucsd.edu under MassIVE ID: MSV000092613. The RNA-sequencing data from nOPCs and aOPCs were previously published and deposited in the Gene Expression Omnibus (GEO) with accession numbers GSE66047 and GSE122446 respectively. ChIP-sequencing data from nOPCs and aOPCs are deposited in GEO with accession number GSE240522.

## ACKNOWLEDGMENTS

This project was supported by R35-NS111604 from the National Institute Neurological Disorders and Stroke of Health to PC. The mass spectrometry experiments are in part supported by a shared instrumentation grant from the NIH, 1S10OD010582-01A1, for the purchase of an Orbitrap Eclipse. We thank the Epigenetics core facility at the Advanced Science Research Center, NY for their help with sorting nOPCs and aOPCs, library preparation for RNA-and ChIP-sequencing and the RNAscope *in situ* hybridization experiments and the NYU Proteomic Core for performing and analyzing unbiased quantitative histone proteomics on our samples. We would like to thank the Casaccia lab members for their constructive feedback during all stages of the project.

## AUTHOR CONTRIBUTIONS

PC worked with D.K.D. on the initial experimental design and conceptualization, wrote the text with D.K.D, and supervised the overall workflow. RNA-sequencing samples were sorted and prepared by S.M. and RNA-sequencing data re-analyzed by D.H. Cell sorting for histone proteomics and ChIP was performed by D.K.D. ChIP was performed by D.K.D. and ChIP-sequencing data analyzed by M.L. Gene ontology analysis was performed by D.K.D. and D.H. Immunohistochemical staining and *in situ* hybridization were performed by D.K.D. and S.S. Western blot analysis, MACS-isolation of OPCs, KATi and SUV420i treatments and RT-qPCR were performed by D.K.D. All the authors provided comments and edited the text.

---

**Supplemental Figure 1.**
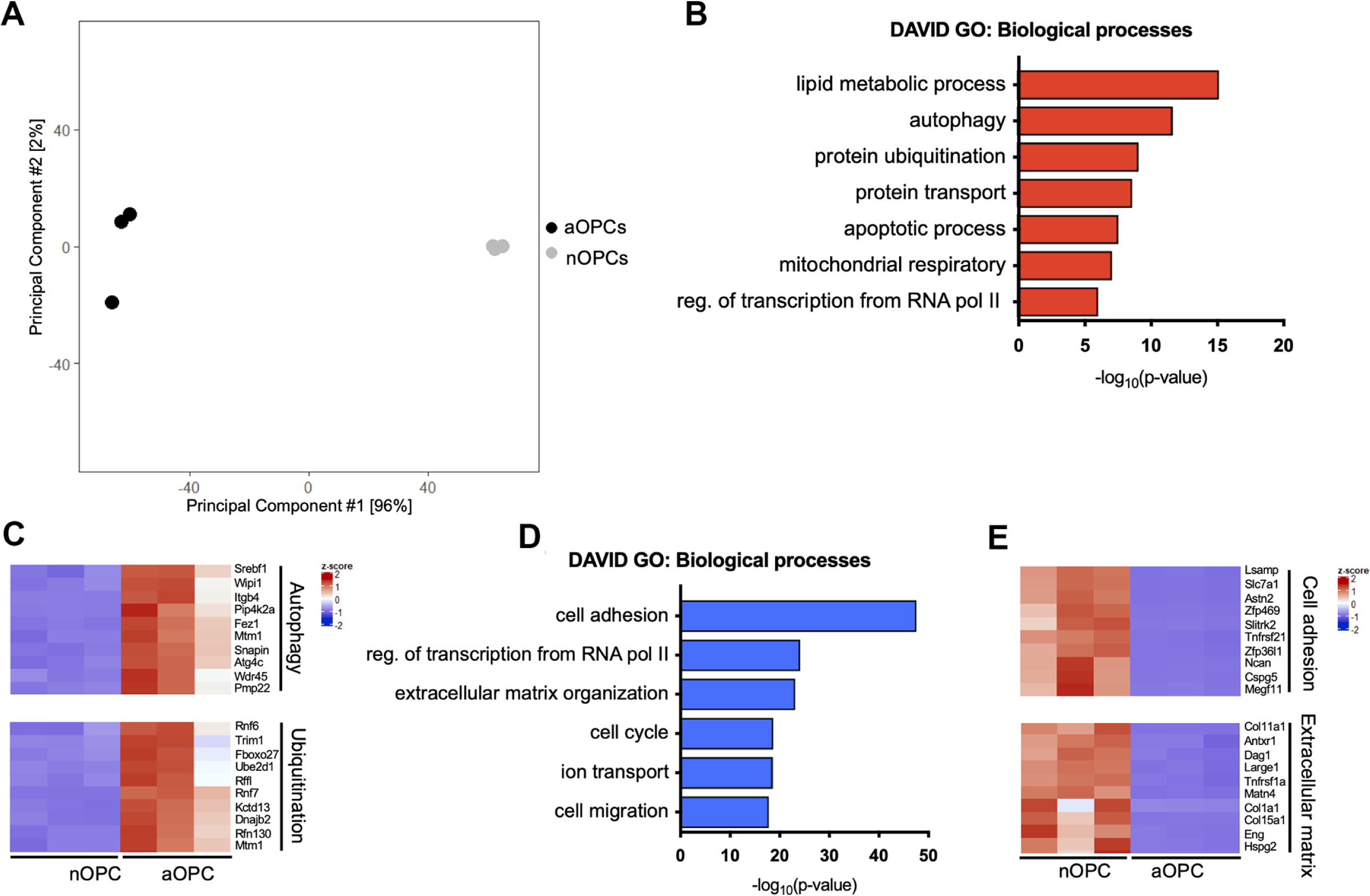
Differential gene expression in aOPCs compared to nOPCs. **(A).** Principal component analysis (PCA) plot assessing the overall quality of RNA-sequencing performed on nOPCs (gray) and aOPCs (black). **(B).** Gene ontology categories related to the 3,406 genes upregulated in aOPCs using DAVID tool. **(C).** Gene expression heatmaps of biological processes related to the upregulated in aOPCs compared to nOPCs. Differentially expressed genes related to autophagy and ubiquitination are shown. **(D).** Ontology categories related to the 5,082 genes downregulated in aOPCs using DAVID tool. **(E).** Gene expression heatmaps of biological processes related to the downregulated in aOPCs compared to nOPCs. Differentially expressed genes related to cell adhesion, and extracellular matrix are shown.

**Supplemental Figure 2.**
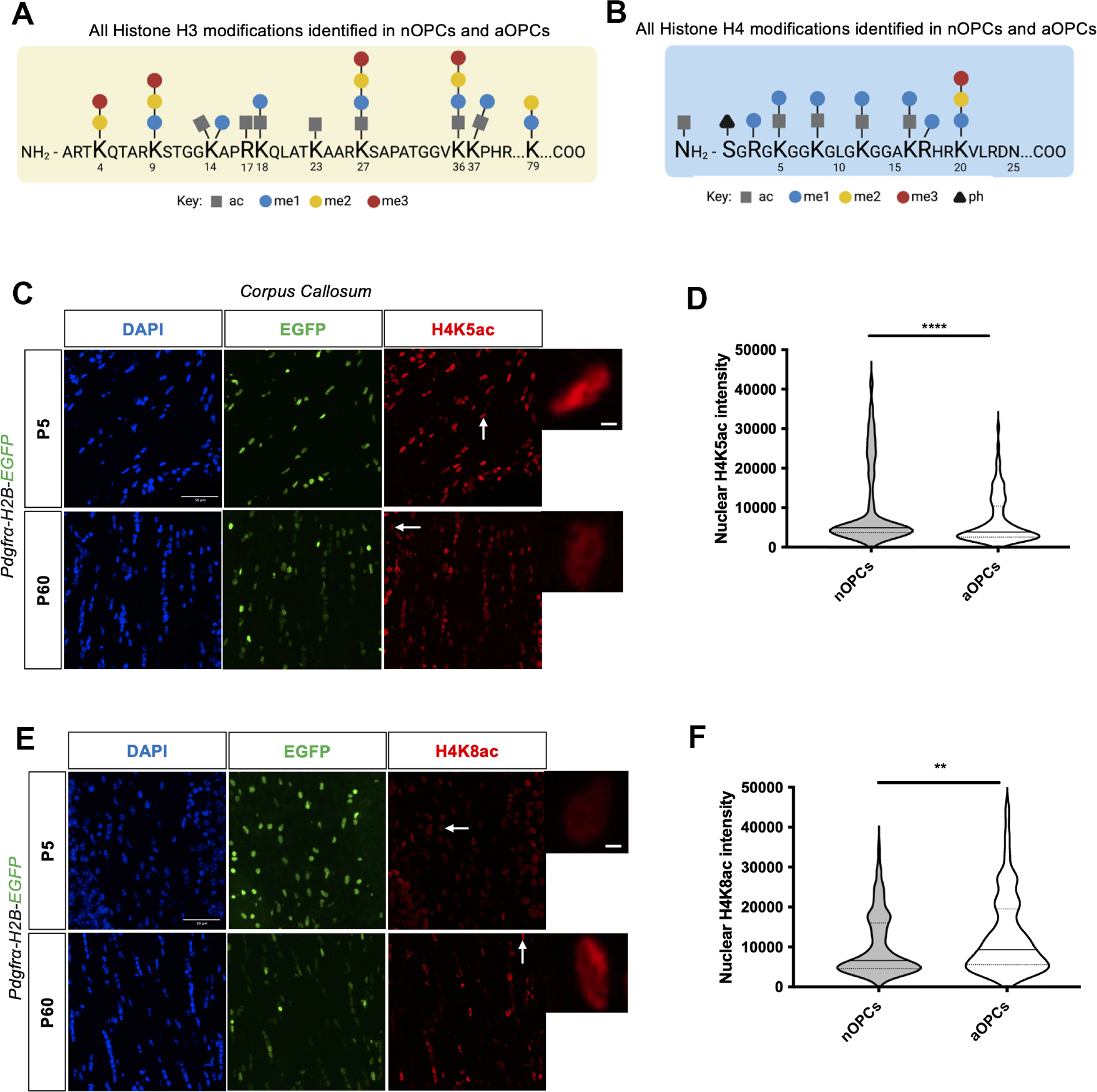
Histone changes in aOPCs compared to nOPCs. Summary of all the histone PTMs identified in nOPCs and aOPCs on **(A)** histone H3 and **(B)** histone H4. **(C).** Immunohistochemical staining of the corpus callosum of *Pdgfra-H2BEGFP* reporter mice with H4K5ac specific antibody (red) in neonatal and adult brain sections. DAPI is stained in blue and PDGFRα^+^ cells are in green. Scale bar = 50 μm. **(D).** Violin plots of nuclear intensity of H4K5ac in PDGFRα^+^ cells in nOPCs and aOPCs. Data represent H4K5ac nuclei intensity of 300 cells from n = 3 mice (****p<0.0001, Student’s *t* test). **(E).** Immunohistochemical staining of the corpus callosum of *Pdgfra-H2BEGFP* reporter mice with H4K8ac specific antibody (red) in neonatal and adult brain sections. **(F).** Violin plots of nuclear intensity of H4K8ac in PDGFRα^+^ cells in nOPCs and aOPCs. Data represent H4K8ac nuclei intensity of 225 cells from n = 3 mice (**p<0.01, Student’s *t* test). DAPI is stained in blue and PDGFRα^+^ cells are in green. Scale bar = 50 μm.

**Supplemental Figure 3.**
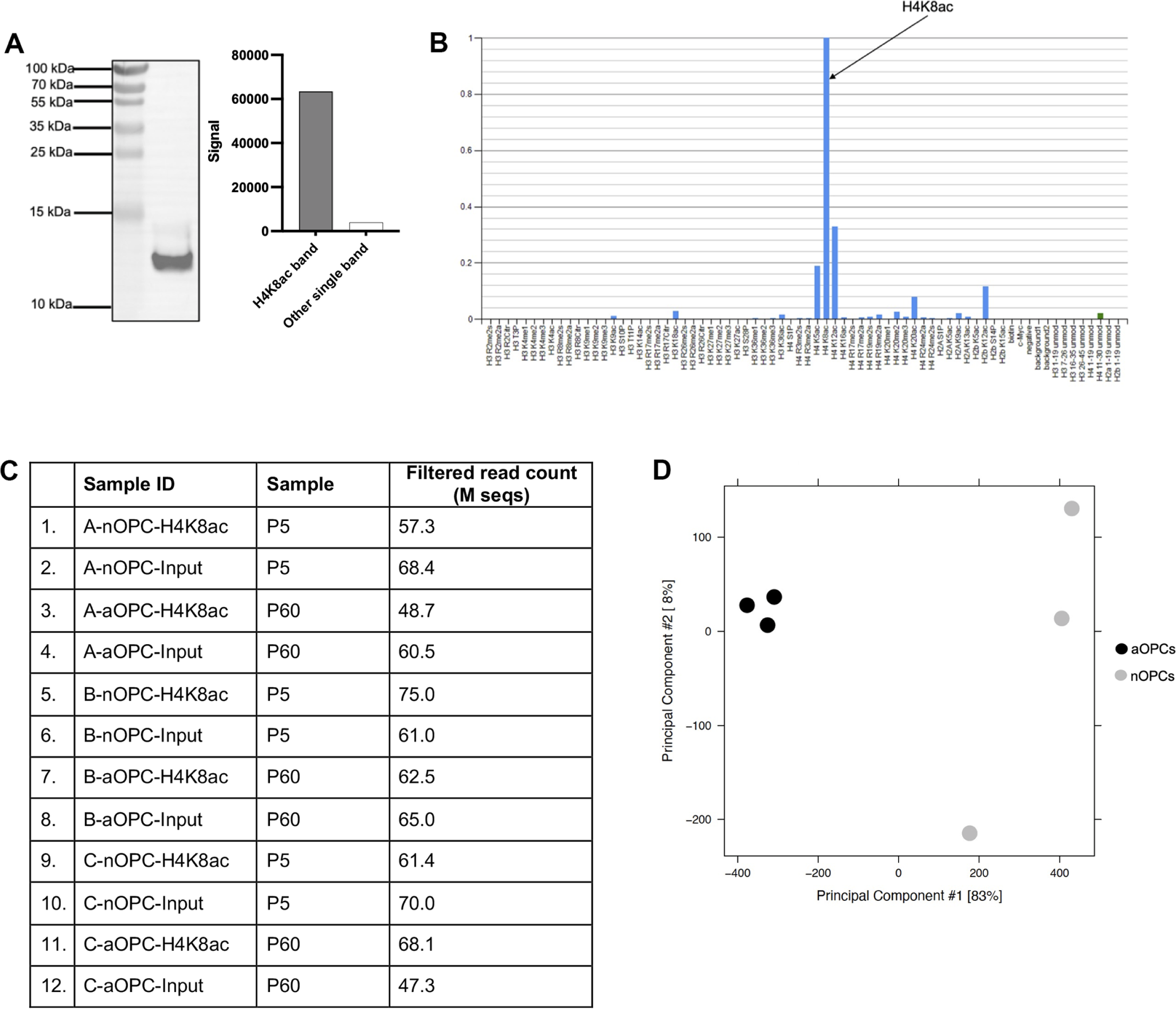
Validation of antibody specificity and overall quality assessment of ChIP-seq data. **(A).** Western blot analysis showing ∼16-fold enrichment of H4K8ac signal relative to any other band in whole cell extracts of oli-neu ceIls (ENCODE guideline - primary characterization). **(B).** Results from the modified peptide array (Active Motif, 13005) used to validate the specificity of the H4K8ac antibody in the presence of competing peptides (ENCODE guideline - secondary characterization). **(C).** ChIP read results for the three immunoprecipitated samples of nOPC and aOPC as well as input chromatin. **(D).** PCA plot assessing the overall quality of ChIP-sequencing performed on nOPCs (gray) and aOPCs (black).

**Supplemental Figure 4.**
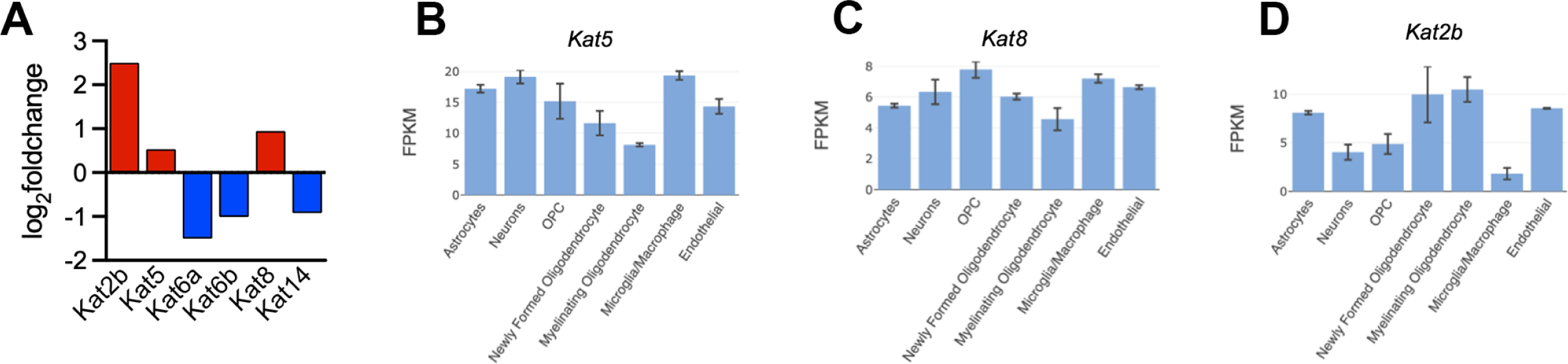
Differential expression of lysine acetyltransferases in oligodendrocyte lineage cells. **(A).** Differential expression of lysine acetyltransferases (KATs) in aOPCs compared to nOPCs. Downregulated KATs and upregulated KATs in aOPCs compared to nOPCs are shown in blue and red respectively. **(B-D).** Transcript levels of *Kat5* **(B)***, Kat8* **(B)** and *Kat2b* **(B)** in different cell types in the brain as reported in previously published RNA-seq database (Zhang et al., 2014).

**Supplemental Table 1.**
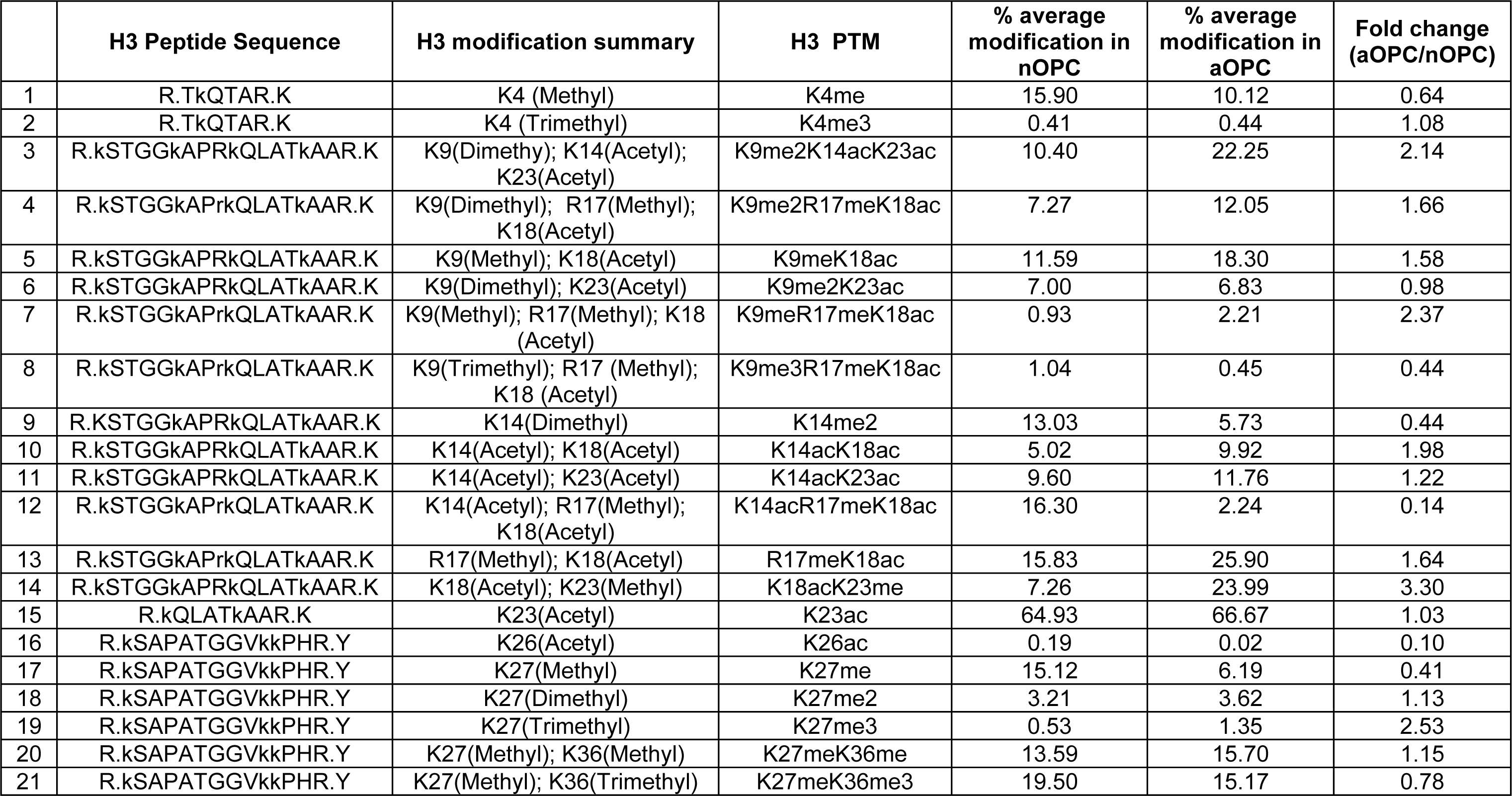
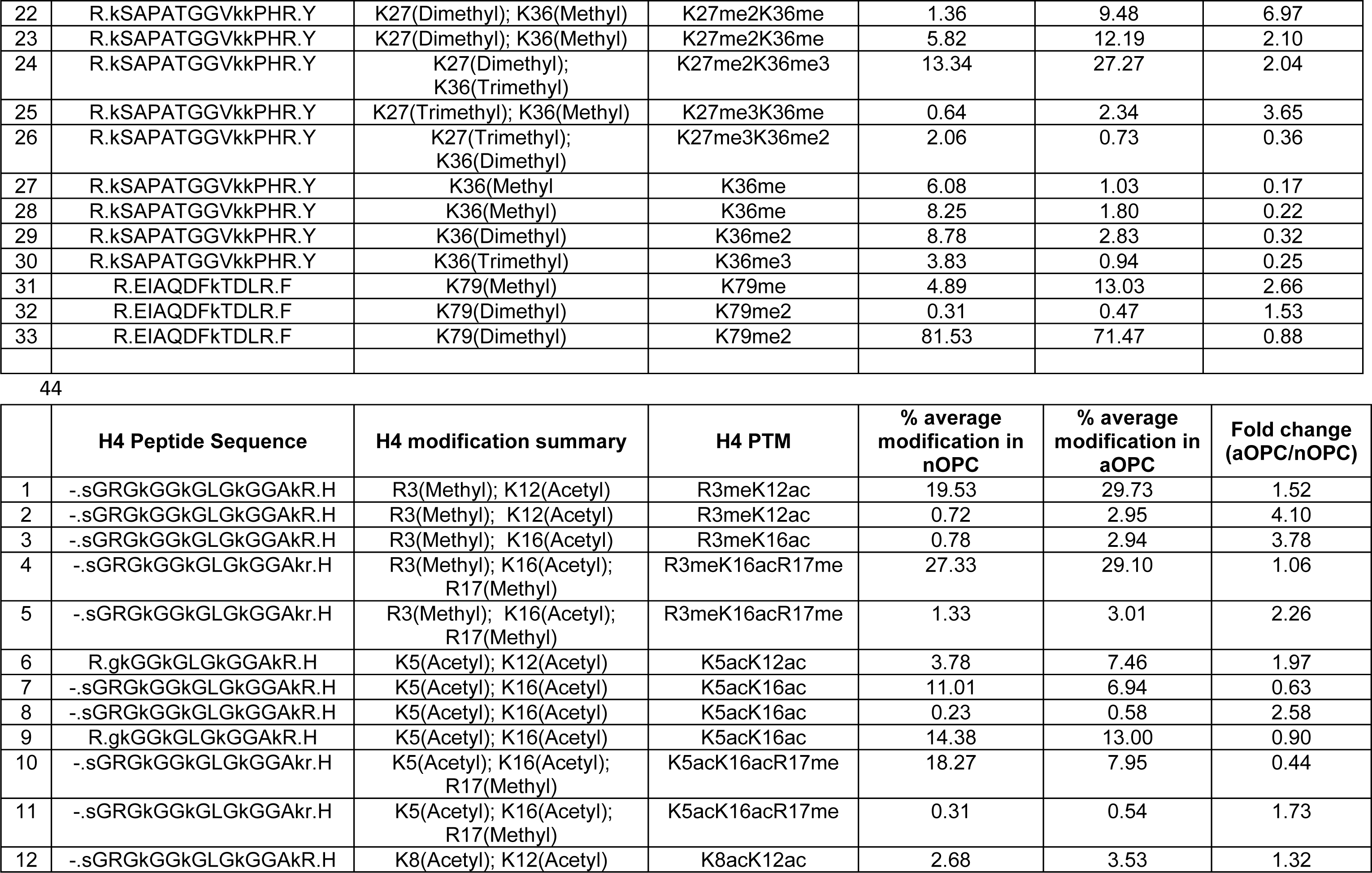
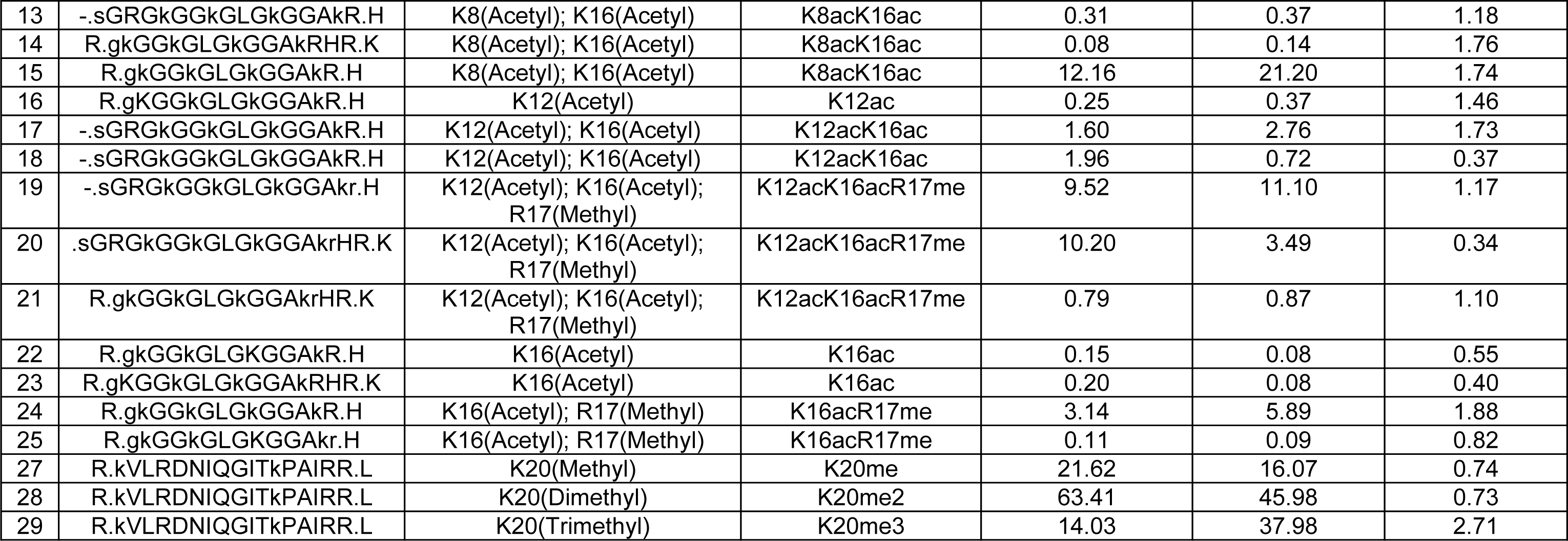
Histone H3 and H4 PTMs identified in oligodendrocyte progenitors. List of all the significant histone PTMs identified on histone H3 and H4 in nOPCs and aOPCs using unbiased quantitative histone proteomics. For each modification, the peptide sequence, modification summary, percent average modification and the ratio between aOPC/nOPC are shown.

